# Statistical and Evolutionary Analysis of Sequenced DNA from Breast Cancer FFPE Specimens

**DOI:** 10.1101/2025.10.04.680485

**Authors:** Monika K. Kurpas, Paweł Kuś, Roman Jaksik, Khanh N. Dinh, Agnieszka Adamczyk, Kaja Majchrzyk, Marek Kimmel

**Affiliations:** Department of Systems Biology and Engineering Silesian University of Technology, Gliwice, Poland; Irving Institute for Cancer Dynamics and Department of Statistics Columbia University, New York, NY, USA; Department of Tumor Pathology Maria Sklodowska-Curie National Research Institute of Oncology, Krakow Branch Krakow, Poland; Departments of Statistics and Bioengineering, and Ken Kennedy Institute Rice University, Houston, TX, USA

**Keywords:** BRCA, cancer, FFPE, molecular evolution

## Abstract

**Background:** Despite the introduction of instant freezing of tumor specimens, formalin-fixed paraffin-embedded (FFPE) blocks of tissue are still commonplace in clinical practice and constitute an important reference for genetic epidemiology of cancer. We carried out a study of a collection of breast tumors paired with lymph-node metastases and analyzed using advanced computational methods, to determine how much information can be obtained from mid-depth whole-exome bulk DNA sequencing.

**Methods:** We gathered 15 paired (primary and an involved lymph node) excised breast tumors of different molecular subtypes (HER2+, triple negative, luminal A and luminal B HER2-), from the National Research Institute of Oncology, Krakow (Poland) Branch. FFPE specimens contained typical artifacts, manifesting themselves in spurious DNA variant calls. We used several bioinformatics tools to remove the artifacts and analyzed the exomic data, using both commercial and original in-house computational techniques.

**Results:** We used several of recent bioinformatics tools to remove the FFPE artifacts and found a serious dispersal of outcomes. After calibration, a series of analyses was performed, including copy number study, resulting in ploidy levels ranging from 1 to 5 (average of 2.5). Positive association was found between the frequency of oncogenes relative to tumor suppressor genes and DNA copy number. In addition, we carried out analyses of the clonal structure of the data using original computational methods based on evolutionary modeling. Interesting results concerning clonal structure, early tumor expansion, and interdependence of the primary tumor and lymph node metastases have been obtained.

**Conclusions:** Despite the imperfections of the FFPE data, many important features of molecular evolution of tumor DNA can be recovered from routine clinical samples.

## 1 Background

The purpose of this paper is to confront the recent ideas of molecular evolution of tumors with DNA-sequencing data obtained from routine pathology specimens of breast cancer, at a major cancer therapy Center in Poland, the Krakow Branch of the Maria Sklodowska-Curie National Research Institute of Oncology. The formalin-fixed, paraffin embedded pathology specimens, we are working with in this paper, are still the work-horse of cancer diagnostics. They may constitute a major resource of regional epidemiology of cancer, if their strengths and weaknesses are investigated and understood.

Breast cancer is one of the most common cancers, and accounts for about one-third of all malignancies in women, with mortality rate about 15% of the total number of cases diagnosed [45]. In 2022, worldwide 2.3 million women were diagnosed with breast cancer and 670,000 women died. Incidence and mortality rate is influenced by age, ethnicity and geography [7]. Worldwide, there were 2.26 million new cases in 2020 almost exclusively in women (over 12% of all new cancer cases and 25% of all new cancer cases in women) [1, 5]. In the same year, it was the cause of nearly 700,000 deaths, making it the fifth most deadly cancer type. In Poland in 2022, according to the Polish National Cancer Registry, the most common cancers among women were breast (23.6%, 21,554 women) and lung (92%) and most deaths were caused by lung cancer (18%) and breast cancer (15%, 6611 women) [3].

Breast cancer is a highly heterogeneous malignancy. The World Health Organization Classification of Breast Tumours (5th Edition) distinguishes 28 subtypes of invasive breast carcinoma. Different breast tumor subtypes are characterized by different risk factors, clinical and histopathological features, outcome, and response to therapies. Histologically the most common invasive type of breast cancer are: invasive ductal carcinoma and invasive lobular carcinoma. Less frequent invasive subtypes include medullary, apocrine, tubular, mucinous, metaplastic, cribriform, neuroendocrine, classic lobular, and pleomorphic breast cancer [7, 19].

Prognostic evaluation, treatment decision making and molecular classification is aided by immunohistochemical staining results for estrogene receptor (ER), progesterone receptor (PR), human epidermal growth factor receptor 2 (HER2) and Ki-67 protein. Four main molecular subtypes are: luminal A, luminal B, HER2-enriched, and triple-negative breast cancer (TNBC) [45, 7, 19]. Luminal A subtype is characterized by ER/PR positivity and HER2 negativity and is most common subtype (68% of cases) [3]. In Luminal A subtype Ki-67 expression is low and this subtype has favorable prognosis and better response to hormone therapy [45, 7, 19]. Luminal B breast cancer expresses the hormone receptor ER and it can be: a) HER2-negative with at least one of: PR low to none (*<*20%) and the biomarker Ki-67 highly expressed (≥ 20%) or b) HER2-positive with variable levels of PR and Ki-67 [7]. Presence of higher levels of Ki-67 than in luminal A, indicates accelerated tumor growth and is associated with worse prognosis [45]. These tumors can be treated with hormone therapy and chemotherapy [45, 7, 19] and encompass about 10% of breast cancers [19]. The HER2-enriched subtype represents 10-20% of breasts cancers, with lack of ER/PR expression and overexpression of HER2; this subtype usually has high expression of Ki-67, is often more aggressive and has a worse prognosis compered to the luminal subtypes [45, 7]. However, patients with HER2 overexpression can benefit from anti-HER2 therapies [45]. Triple negative subtype does not express ER, PR, or HER2 and represents 10-20% of breast cancers [45, 7, 19]. TBNC is the most aggressive of the four subtypes, with the highest proliferation index of Ki-67, with higher risk of recurrence and poorer prognosis [45, 7]. TNBC often harbors DNA alterations and genetic mutations, mostly associated with mutation in the BRCA1/2 genes [45, 7]. TNBC is also called basal-like, characterized by the absence of all 3 receptors.

Breast cancer (BRCA) originates from the epithelial cells and show high mRNA signatures of Epithelial–Mesenchymal Transition (EMT) [21], a mechanism that may support cancers metastasis. To study the evolution and metastasis of BRCA, we performed the whole exome sequencing of the data from 15 BRCA patients, using paired primary tumor and invaded lymph node (denoted P1 and L1, respectively), as depicted in Tables 1 and 2.

**Table 1:**
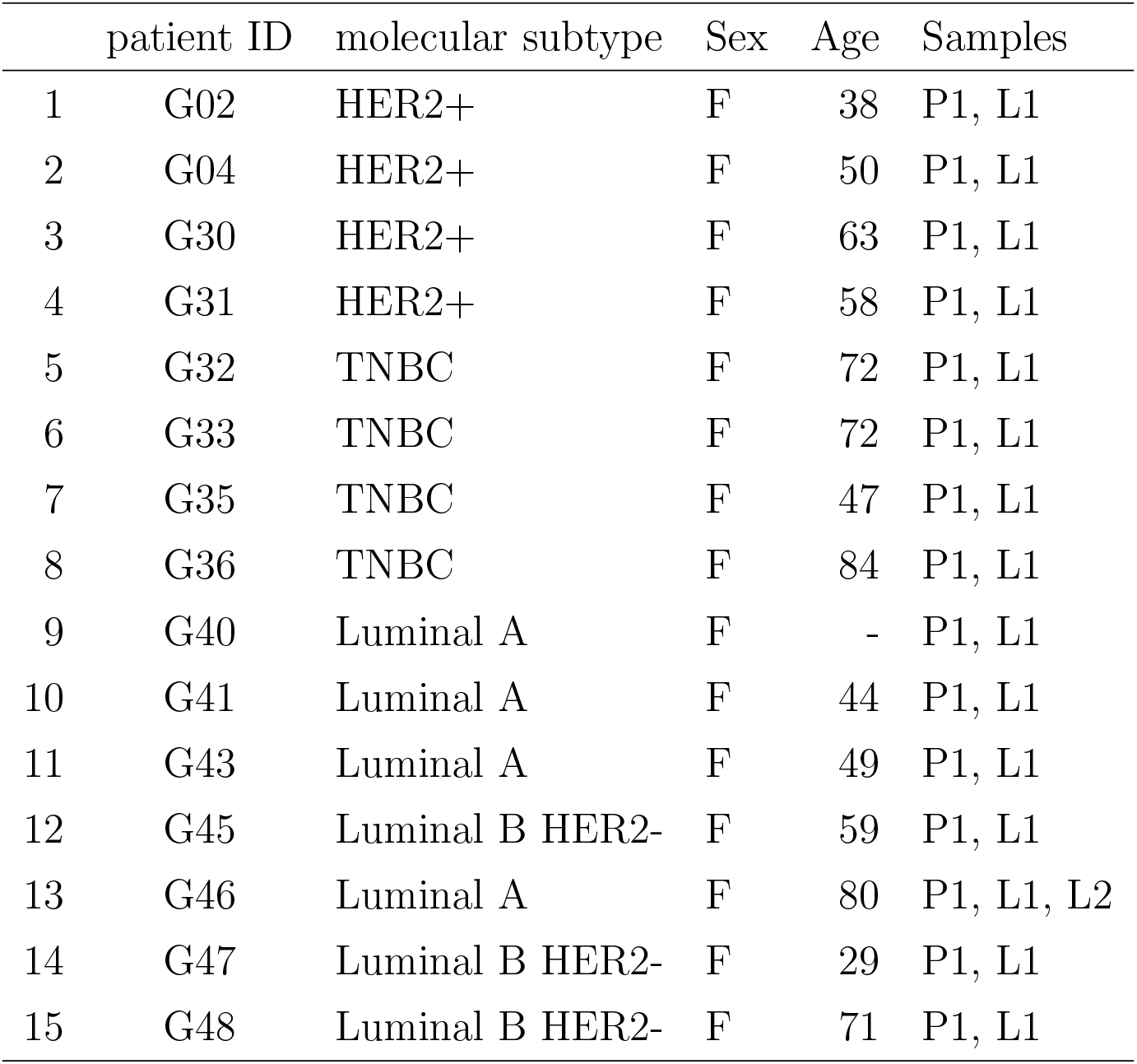
List of BRCA patients. Two tumor samples were obtained and sequenced from each patient. *P1* primary tumor sample, *L1* - lymph node metastasis sample, *F* - female. In the G46 case, two lymph nodes were ascertained.

|  | patient ID | molecular subtype | Sex | Age | Samples |
| --- | --- | --- | --- | --- | --- |
| 1 | G02 | HER2+ | F | 38 | P1, L1 |
| 2 | G04 | HER2+ | F | 50 | P1, L1 |
| 3 | G30 | HER2+ | F | 63 | P1, L1 |
| 4 | G31 | HER2+ | F | 58 | P1, L1 |
| 5 | G32 | TNBC | F | 72 | P1, L1 |
| 6 | G33 | TNBC | F | 72 | P1, L1 |
| 7 | G35 | TNBC | F | 47 | P1, L1 |
| 8 | G36 | TNBC | F | 84 | P1, L1 |
| 9 | G40 | Luminal A | F | - | P1, L1 |
| 10 | G41 | Luminal A | F | 44 | P1, L1 |
| 11 | G43 | Luminal A | F | 49 | P1, L1 |
| 12 | G45 | Luminal B HER2- | F | 59 | P1, L1 |
| 13 | G46 | Luminal A | F | 80 | P1, L1, L2 |
| 14 | G47 | Luminal B HER2- | F | 29 | P1, L1 |
| 15 | G48 | Luminal B HER2- | F | 71 | P1, L1 |

**Table 2:** ASCAT estimates of tumor purity and ploidy in BRCA cohort. *P1* - primary tumor sample, *L1* lymph node metastasis sample. G43 L1 sample’s estimate is not available. In the G46 case, two lymph nodes were ascertained.

| Patient ID | Sample | Purity | Ploidy | Molecular subtype |
| --- | --- | --- | --- | --- |
| G02 | P1 | 0.23 | 3.78 | HER2+ |
| G02 | L1 | 1 | 2.02 | HER2+ |
| G04 | P1 | 0.58 | 1.56 | HER2+ |
| G04 | L1 | 0.31 | 4.52 | HER2+ |
| G30 | P1 | 0.43 | 3.06 | HER2+ |
| G30 | L1 | 0.48 | 3.58 | HER2+ |
| G31 | P1 | 0.35 | 3.00 | HER2+ |
| G31 | L1 | 0.5 | 2.86 | HER2+ |
| G32 | P1 | 0.47 | 1.86 | TNBC |
| G32 | L1 | 0.43 | 3.44 | TNBC |
| G33 | P1 | 0.59 | 1.65 | TNBC |
| G33 | L1 | 0.62 | 1.55 | TNBC |
| G35 | P1 | 0.66 | 1.95 | TNBC |
| G35 | L1 | 0.44 | 1.89 | TNBC |
| G36 | P1 | 0.61 | 3.16 | TNBC |
| G36 | L1 | 0.31 | 3.21 | TNBC |
| G40 | P1 | 0.75 | 2.09 | Luminal A |
| G40 | L1 | 0.89 | 2.22 | Luminal A |
| G41 | P1 | 0.39 | 2.06 | Luminal A |
| G41 | L1 | 1 | 2.01 | Luminal A |
| G43 | P1 | 0.61 | 2.12 | Luminal A |
| G45 | P1 | 0.71 | 3.17 | Luminal B HER2- |
| G45 | L1 | 0.48 | 3.40 | Luminal B HER2- |
| G46 | P1 | 0.82 | 2.18 | Luminal A |
| G46 | L1 | 0.56 | 2.39 | Luminal A |
| G46 | L2 | 0.69 | 2.38 | Luminal A |
| G47 | P1 | 0.58 | 1.82 | Luminal B HER2- |
| G47 | L1 | 0.63 | 1.84 | Luminal B HER2- |
| G48 | P1 | 0.62 | 2.10 | Luminal B HER2- |
| G48 | L1 | 0.54 | 2.09 | Luminal B HER2- |

The FFPE specimens contained artifacts typical for the fixation and preservation method (see further on), resulting in spurious DNA variant calls. To remove these calls, we used the bioinformatics tools, SOBDetector [14], FFPolish [16], DEEPOMICS and DEEPOMICS plus [26], which identify context-dependent substitution signatures, but also other features such as variant frequency and strand directionality. We found discrepancies in outcomes between the tools.

Aside from these important technical questions, our major purpose was to employ the variant allele frequencies (VAF) synonymous with the site frequency spectra (SFS) to gather conclusions concerning the clonal evolution of the breast cancer specimens. For this purpose, we employed decomposition of the spectra developed by us in [11]. The analysis resulted in interesting conclusions concerning clonal expansion and mutation and growth rates.

In addition, we carried out copy number analysis, resulting in ploidy levels ranging from 1 to 5 (average of 2.5). One of the reasons to perform the analysis was to verify if a positive association between the DNA copy number and frequency of oncogenes relative to tumor suppressor genes [43] and other evolutionary characteristics holds in our data.

## 2 Methods

### 2.1 DNA sequencing of cell samples from breast cancer specimens

#### 2.1.1 DNA sample collection and processing

Paired tissue samples from primary breast tumor locations and concurrent metastases to regional lymph nodes, along with control samples from not involved lymph node, were collected in the Department of Tumor Pathology of the Maria Sklodowska-Curie National Research Institute of Oncology, Krakow (Poland) Branch. Cancer specimens were matched with normal tissue samples used as a reference for individual genetic background (control samples). Of all 3-sample sets from 15 patients, four were HER2+ breast cancer subtypes (specimens G2, G4, G30 and G31), four were triple-negative (specimens G32, G33, G35 and G36), four were luminal A subtype (specimens G40, G41, G43 and G46) and remaining three were luminal B HER2-subtype (see Table 1) according to classification in [22] and thresholds provided in [7].

DNA samples were isolated in the Department of Tumor Pathology from macrodissected FFPE tissue samples using a semi-automatic method available from Maxwell^®^ RSC Instrument (Promega) with the settings recommended by the manufacturer. The quality and amount of DNA was evaluated using the NanoDrop 2000c spectrophotometer and fluorimetrically using the Qubit™ dsDNA HS Assay Kit and the Qubit 3.0 device (ThermoFisher Scientific). Genomic DNA from each sample was subjected to library preparation using the Agilent SureSelect Human All Exon V6 kit. Sequencing was carried out using the Illumina NovaSeq X platform in a paired-end configuration with 150bp reads. Each sample achieved a throughput of approximately 15Gbp. Both library preparation and sequencing was performed by Novogene Europe.

#### 2.1.2 DNA sequencing Quality Control

Quality control of raw sequencing reads was performed using FastQC and FastQ Screen. Reads were aligned to the GRCh38 reference genome using BWA-MEM (v0.7.17) [30], with support for alternative contigs enabled. Post-alignment processing included duplicate marking using the MarkDuplicates algorithm from the Picard toolkit and base quality score recalibration with BaseRecalibrator, a component of the Genome Analysis Toolkit (GATK v4.2.6.1) [10]. Somatic variants were identified using MuTect2 (v4.2.6.1) [10] based on matched tumor-normal sample pairs.

Shown below are the representative examples of coverage charts and read statistics (Figure 1 and Figure 2, respectively) for samples G30 and G31. The coverage histograms and the read statistics for all samples are in Figs S1 and S2. The sequencing statistics are listed in Supplementary Table 1. The read statistics for all samples are subdivided into primary tumor (P1), cancerous lymph node (L1), and not involved lymph node (C), this latter providing control for detection of somatic mutations. Based on our experience the quality of data obtained based on the FFPE material is comparable to the quality of data obtained typically using fresh frozen (FF) material. We were able to obtain *>*100x true median coverage for the regions captured by the Agilent SureSelect Human All Exon v7 probes, as shown on Figure 1. This was achieved both due to high quality of the reads, moderate read duplication rate for the total number of reads sequenced (*>*100 M/sample) and high mappability rate, as shown on Figure 2.

**Figure 1:**
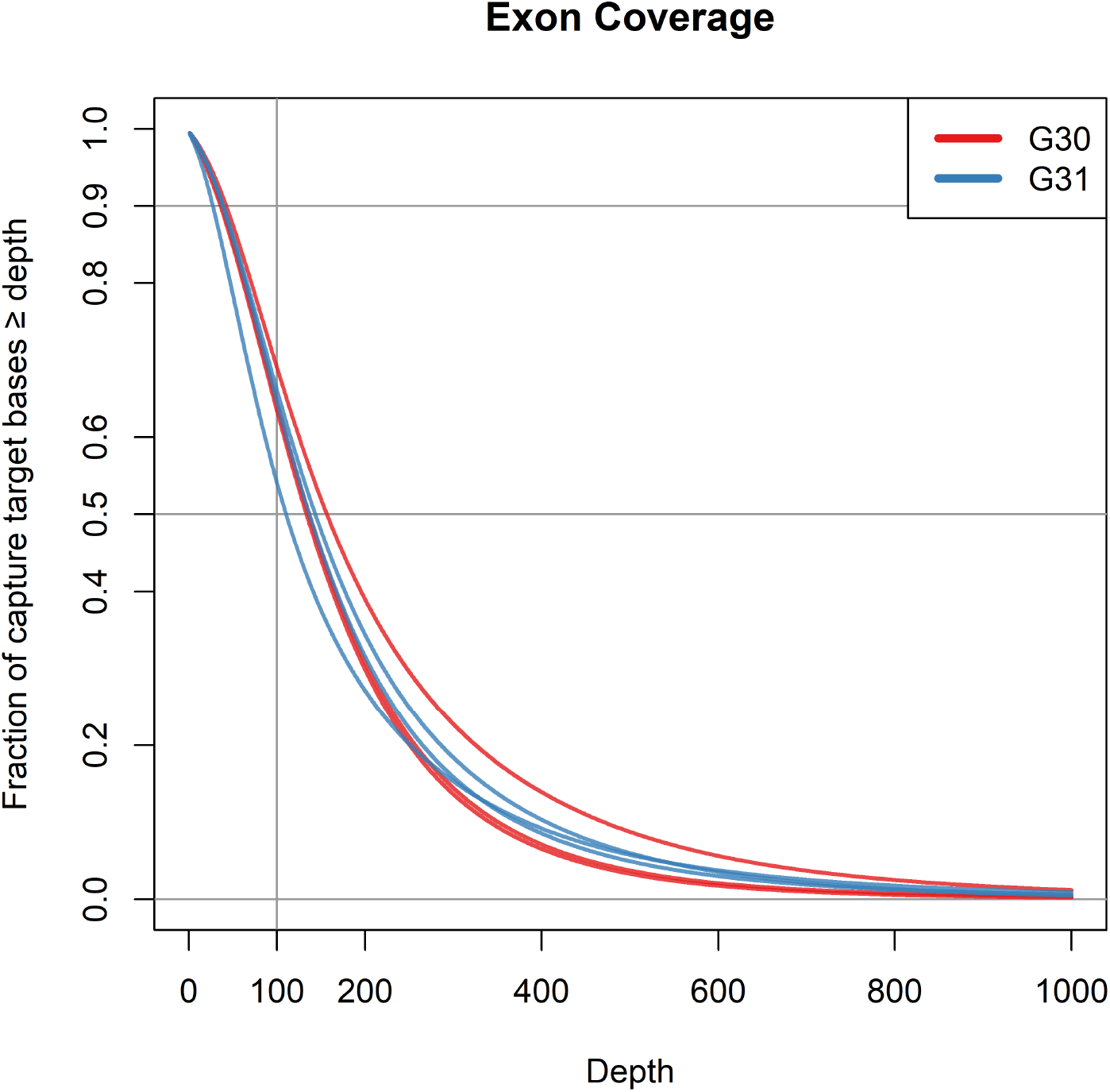
Exon coverage characteristics for the G30 (red) and G31 (blue) patient samples.

**Figure 2:**
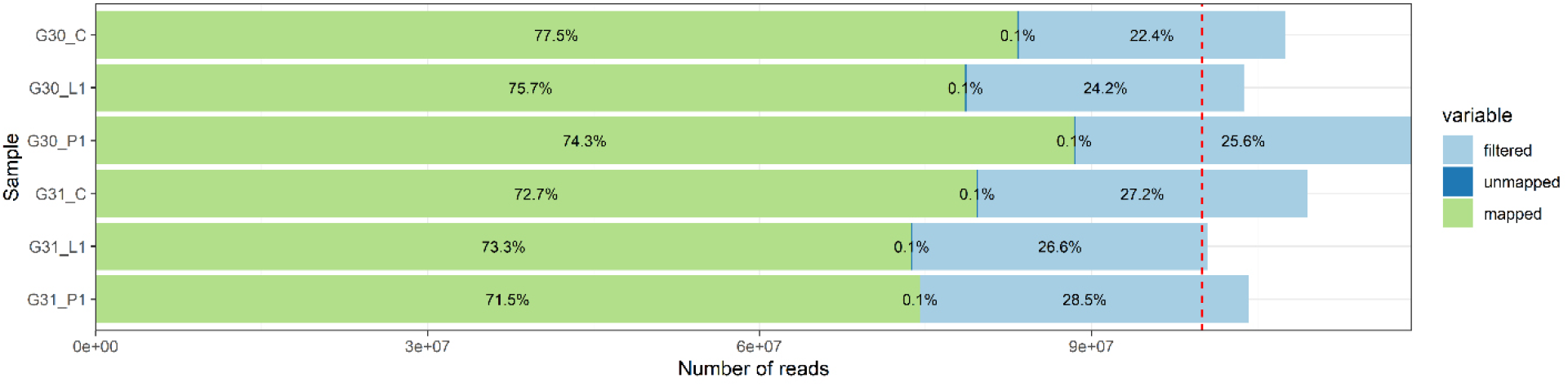
Reads statistics for G30 and G31 patient samples.

While we are unable to compare the results based on FFPE and FF samples obtained from the same individuals it seems safe to assume that for isolated DNA of good quality the results can be comparable between FFPE and FF, as shown in [6].

Variant filtering was carried out with GATK’s FilterMutectCalls, incorporating sample contamination estimates obtained via the CalculateContamination tool and read orientation bias metrics derived from the LearnReadOrientationModel tool. All high-confidence variants were annotated using the Variant Effect Predictor (VEP v107) [33].

#### 2.1.3 Removal of formalin fixation artifacts

Formalin-fixed, paraffin-embedded (FFPE) samples are among the most common type of material collected in clinical settings due to the short time of sample preparation and the high durability of preserved blocks [16]. They are frequently used as specimens for histopathological and molecular studies. However, the method of fixation with formalin causes nucleic acid damage which poses challenge to using it as a source of genetic material for sequencing experiments. The artifacts of the fixation with formalin (called further FFPE artifacts) arise mainly by deamination of cytosine to uracil, which results in mismatched pairing by DNA polymerases [15]. In the sequencing data, this is seen as enrichment in number of C*>*T and G*>*A (the opposite strand) substitutions and overall a considerable number of spurious mutational variants compared to the fresh-frozen samples [15]. To account for the necessity of eliminating false-positive mutational variants, several strategies has been developed, including enzymatic methods of DNA repair during isolation of genetic material from FFPE blocks [8] and a set of bioinformatic approaches reviewed and compared with reference data in [37].

Due to the fact that the genetic material was already sequenced, we did not have the possibility to chemically remove FFPE artifacts which appears to be the most reliable method [37]. Instead we need to rely on bioinformatic tools. In this work four of them were applied and their results tested: SOBDetector[14], FFPolish[16], DEEPOMICS [26] and DEEPOMICS plus (unpublished release of the same group). For the details regarding the tools characteristics, methods of working, input files and achieved efficacy see [37].

Whole exome sequencing data described in this work were prepared accordingly to the requirements of the tools. SOBDetector (v1.0.4) was executed using the original VCF and BAM files for each sample. The tool annotated variants with an artiStatus label in the VCF INFO field. Variants were labeled as “artifact”, “snv” and “uncategorized”. FFPolish (v0.1) was executed using the reference genome and sample-specific BAM files. For each sample, the tool produced a filtered VCF file with FFPE-induced artifacts removed. Additional annotations were added to indicate whether variants were retained or removed during filtration. DEEPOMICS FFPE (and DEEPOMICS FFPE plus) analysis was conducted using the on-premise version of the tool, in collaboration with Theragen Bio. The same software is also accessible via the web portal at [2]. The pattern of variants retained by the tools listed above is shown in Figure S4. The numbers were highest in case of SOBDetector, significantly exceeding these observed after filtration with other tools and also the average number of variants observed in The Cancer Genome Atlas (TCGA) [4] samples (see Results, section 3.1.1). In turn, the counts od variants obtained with other tools were too low for some sorts of analyses we conducted (see later on), thus we decided to manipulate the FFPE score threshold in DEEPOMICS result achieving more variants with lower allele frequency but still satisfactory efficacy of artifact filtration (see Results, section 3.1.2).

#### 2.1.4 Ploidy and purity estimates

Purity and ploidy estimates were obtained using ASCAT (Allele-Specific Copy Number Analysis of Tumors) [42] which is an R package that infers tumour purity, ploidy and allele-specific copynumber profiles from LogR and BAF tracks derived from the sequencing reads using tumor/normal BAM pairs. It provides preprocessing (logR correction and germline-genotype prediction), allele-specific segmentation and a purity–ploidy fitting procedure for robust copy-number calling in tumor samples. Table 2 shows the purity and ploidy of the samples estimated by ASCAT. We calculated baseline ploidy of the sample based on weighted average of copy number estimated by ASCAT in genomic regions of a given sample. In some analyses, we used also the CNV levels of specific genome segments.

For part of the samples we also obtained histopathology estimates of sample purity ranging from 80 to 95 percent. Because the values seem to be unrealistically high for breast cancer [25] and that we do not have these estimates from all samples, we decided to use ASCAT purity estimates instead.

#### 2.1.5 Mutational signatures

We used the R package mutSignatures together with BSgenome.Hsapiens.UCSC.hg38 to annotate each single-base substitution with its genomic (trinucleotide) context against the hg38 reference and to build the mutation-count matrix. Then, we used SigProfiler to perform mutational-signature analysis on that matrix, extracting and fitting mutational signatures and matching them to known COSMIC signatures.

### 2.2 Data and statistical analysis

#### 2.2.1 Site frequency spectra

As mentioned in the Introduction, in a sequencing experiment with a known number of sequences we can estimate, for each site at which a novel somatic mutation has arisen, the number of cells carrying that mutation. Inference from evolutionary models of DNA often exploits summary statistics of the sequence data; a common one is the so-called site frequency spectrum. Figure 3 gives an example: time runs down the page.

**Figure 3:**
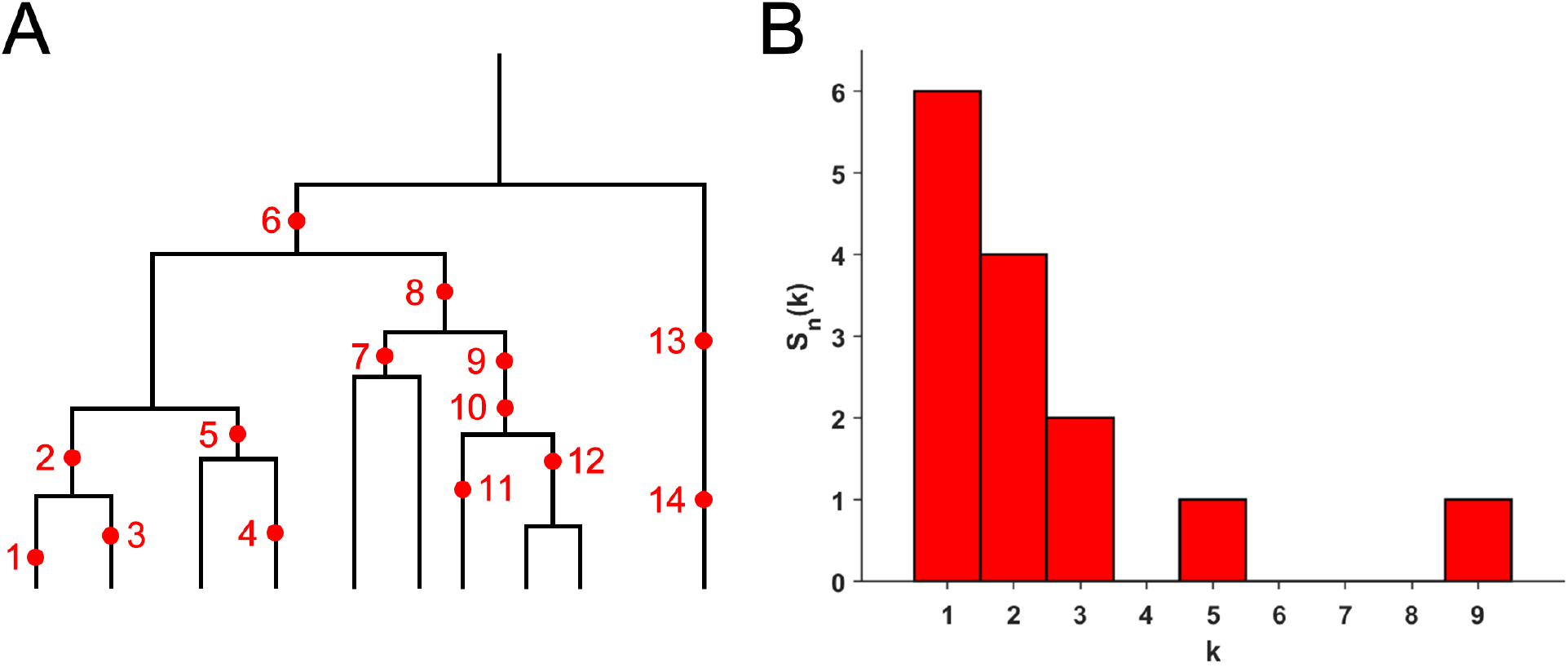
**(A)** The genealogy of a sample of *n* = 10 cells includes 14 mutational events. Mutations 1, 3, 4, 11, 13 and 14 (six mutations) are present in a single cell each; mutations 2, 5, 7 and 12 (four mutations) are present in two cells; mutations 9 and 10 (two mutations) are present in three cells; mutation 8 is present in five cells and mutation 6 is present in nine cells. **(B)** The observed site frequency spectrum, *S*_14_(1) = 6, *S*_14_(2) = 4, *S*_14_(3) = 2, *S*_14_(5) = 1 and *S*_14_(9) = 1, other *S*_*n*_(*k*) equal to 0.

The genealogy of a sample of *n* = 10 cells includes 14 mutational events. Mutations 1, 3, 4, 11, 13 and 14 (six mutations) are present in a single cell each; mutations 2, 5, 7 and 12 (four mutations) are present in two cells; mutations 9 and 10 (two mutations) are present in three cells; mutation 8 is present in five cells and mutation 6 is present in nine cells. If we denote the number of mutations present in *k* cells by *S*_*n*_(*k*), then in this example *S*_*n*_(1) = 6, *S*_*n*_(2) = 4, *S*_*n*_(3) = 2, *S*_*n*_(5) = 1, and *S*_*n*_(9) = 1, with all other *S*_*n*_(*j*) = 0. The vector (*S*_*n*_(1), *S*_*n*_(2), …, *S*_*n*_(*n* − 1)) is called the (observed) site frequency spectrum, abbreviated to SFS.

By convention, one includes only sites that are segregating in the sample — that is, sites at which both the mutant and ancestral types are observed among the sampled cells. Mutations that occurred before the most recent common ancestor of the sampled cells are present in every cell in the sample; these non-segregating mutations are referred to as truncal mutations.

In most cancer sequencing experiments, we do not know the number of cells that were sampled, and, indeed, the DNA sequence of each cell cannot be determined from bulk sequencing data. Nonetheless, we can estimate the relative proportion of the mutant at each segregating site, and so arrive at a frequency spectrum based on proportions. Accordingly, instead of writing *S*_*n*_(*k*), we write *S*(*x*) = *S*(*k/n*), with *x* treated as a continuous variable, such that *x* ∈ (0, 1). We continue to use the term SFS for such a spectrum, as there should be no cause for confusion. In essence, *S*(*x*) is an idealized version of the empirical variant allele frequency (VAF) graph. In addition, it is convenient for reasons explained in Section 2.4 of [27] to define the cumulative tail of the SFS *S*(*x*)

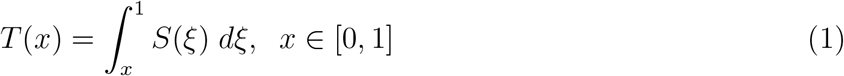

The theory that allows computing the expectations of SFS in populations with a given growth law under the Infinite Site Model (ISM) of mutation, was developed concurrently by many researchers, with one of the seminal papers published in 1998 by Griffiths and Tavaré [23]. The Griffiths-Tavaré expressions are accurate but quite complicated. A computational method which works fast even with very large sample sizes, was developed in a series of papers by Polanski, Kimmel, and co-workers [38]. Tractable approximations were derived under the exponential growth hypothesis by Durrett [18]. A related approach based on linear birth-and-death processes is that by Lambert and co-workers [28].

#### 2.2.2 Variant allele frequency (VAF), site frequency spectra (SFS), and cancer cell fraction (CCF)

This section is based on Reference [9] with slightly modified notation. The basic statistic that DNA sequencing provides for each mutation is the Variant Allele Frequency (VAF):

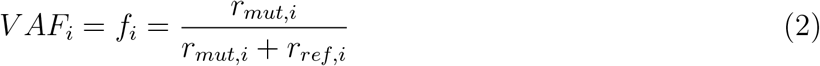

where *r*_*mut*,*i*_ and *r*_*ref*,*i*_ are the numbers of reads supporting the mutant (alternate) and reference alleles, respectively. The convention we accept in this paper is that VAF is the observed manifestation of the idealized SFS introdueced in the previous section. The sum of *r*_*mut*,*i*_ and *r*_*ref*,*i*_ is the depth of variant sequencing:

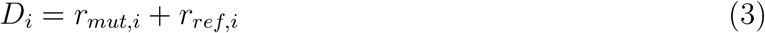

In addition to this it is useful to represent the propensity of mutations or mutation clusters through their “cellular prevalence” (CP, the fraction of cells carrying the mutation or mutations in the sample), or their “cancer cell fraction” (CCF, the fraction of tumor cells carrying the mutation or mutations).

The copy number state *m*_*i*_ of an SNV, also called its “multiplicity”, is key to understanding VAF distributions of mutations. It is helpful to consider the product of mutation multiplicity *m*_*i*_ of a mutation *i* and its cancer cell fraction CCF_*i*_:

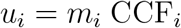

We can relate *u*_*i*_ and *m*_*i*_ by the expression

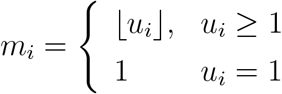

where

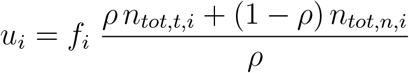

In the expression above, *ρ* and *n*_*tot*,*t*,*i*_ can be obtained through copy number analysis, *f*_*i*_ can be calculated from *r*_*mut*,*i*_ and *r*_*ref*,*i*_ using Equation 1, and the *n*_*tot*,*n*,*i*_ values are considered known (typically equal to 2).

If we consider the bulk DNA sequencing in terms of drawing random sequences from the pool of DNA extracted from the wildtype and mutated cells, *r*_*mut*,*i*_ can be described by the binomial distribution:

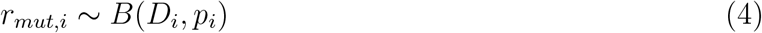

where *p*_*i*_ is the frequency of the mutated allele in the sample, estimated by

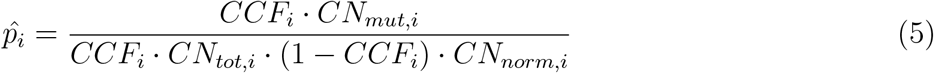

where *CN*_*mut*,*i*_ is the number of copies of the mutant allele in the mutant cells, *CN*_*tot*,*i*_ is the total copy number in the mutated cells, and *CN*_*norm*,*i*_ is the ploidy of the normal cells. In the purely diploid population, where *CN*_*mut*,*i*_ = 1, and *CN*_*tot*,*i*_ = *CN*_*norm*,*i*_ = 2, *p* equals:

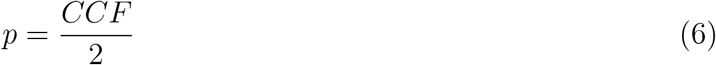

and

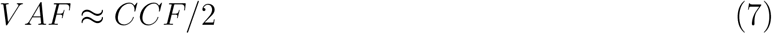

Because of this, due to the complexity associated with the estimation of allele-specific copy numbers, many methods use VAF as an approximate measure of the CCF. In particular, VAF spectra, which represent the distributions of observed allelic frequencies for all variants in a sample, have been found useful in the modeling of neutrality and selection in cancer research.

#### 2.2.3 Inferring parameters for the neutral tail and mutation clusters from observed site frequency spectra

Assuming a tumor consisting of *H* subclones, the expected number of mutations present in *k* copies among a total of *n* alleles in the sequencing sample, following population genetics approaches such as [23, 17, 11], is:

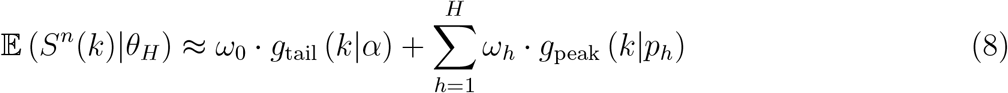

where *ω*_0_ is the number of mutations occurring neutrally in each clone, which follow a power tail distribution *g*_tail_ (*k*|*α*) ∼ 1*/k*^*α*^ with power *α*. Each clone *h* expands from a most recent common ancestor (MRCA) that already contained *ω*_*h*_ mutations, which are inherited by the clone’s progeny. These mutations follow 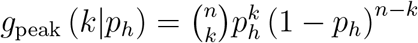, where *p*_*h*_ is the clone’s mean variant allele frequency (VAF). Therefore, the true site frequency spectrum (SFS, Eq. 8) is characterized by parameter set *θ*_*H*_ = {*ω*_0_, *α, ω*_1_, *p*_1_, …, *ω*_*H*_, *p*_*H*_ }.

In our earlier work, we derived a sampling formula for the observed SFS that incorporates sequencing coverage [11], and pertinent details are recounted here. We define *φ*_*r*_ to be the probability that a mutated site is sequenced with *r* reads. We assume these reads originate from distinct cells, as the mean sequencing coverage (≈ 100) is magnitudes lower than the cell count (≈ 10^5^ − 10^6^) in sequencing samples. Given a mutation present in *k* copies and sequenced with *r* reads, each read has probability *k/n* to exhibit the mutation, therefore the alternative read count follows *z*|*r* ∼ Binomial(*r, k/n*). The expected number of mutations with observed VAF *f* ∈ (*f*_1_, *f*_2_], for 0 ≤ *f*_1_ *< f*_2_ ≤ 1, then follows:

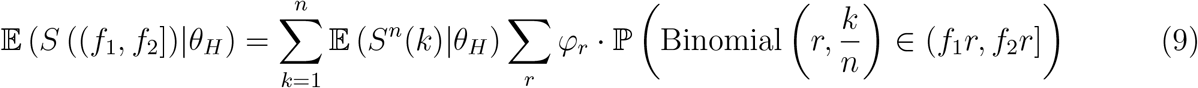

For the computations, we assume sequencing samples with *n* = 1000 copies. The observed SFS 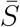 is tabulated from data with bin size *b* = 0.01. We then seek the parameter set *θ*_*H*_ such that Eq. 9 best matches 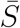 for each bin (*f*_1_, *f*_2_] = (0, *b*], (*b*, 2*b*], …, (1 − *b*, 1].

To find *θ*_*H*_, we employ Approximate Bayesian Computation sequential Monte Carlo with random forests (ABC-SMC-DRF), a method we previously developed for likelihood-free parameter inference [12]. We assume prior distributions *α* ∼ Uniform(0.5, 5), *p*_*h*_ ∼ Uniform(0, 1) and Uniform 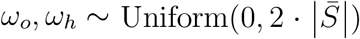, where 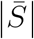 is the number of mutations in the data. ABC-SMC-DRF samples parameter sets *θ*_*H*_ from these priors, computes the corresponding expected SFS from Eq. 9, then evaluates the error 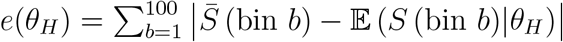. A random forest is contructed from the combined reference table, and the posterior distribution for *θ*_*H*_ is extracted by applying target statistic *e*(*θ*_*H*_) = 0. ABC-SMC-DRF performs this step iteratively, where the parameter sets in later iterations are drawn from the posterior distribution of the previous step. The result from the final iteration contains the joint posterior distribution for *θ*_*H*_. We will skip further details since they can be found in a forthcoming publication.

#### 2.2.4 Deterministic models of SFS under neutrality and population growth

In this section, we show a simple theory, which allows interpreting SFS power function tail components of the SFS with exponents different from −2. Spectra with power exponent equal to −2 were previously used in the intuitive MOBSTER software by Williams and Co-workers [44] and are believed to represent the neutrally evolving subpopulations of exponentially-growing cell populations with constant mutation rate. The area *A* under this component of the spectrum can be proved to equal *n* times the “reduced mutation rate” *µ/r* (roughly, mutation rate per cell division), where *n* is sample size equivalent to the sequencing depth and *µ* and *r* are rates of mutation and exponential growth respectively. This allows us to estimate the reduced mutation rate by

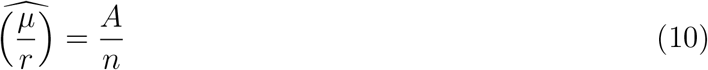

Durrett, in reference [17], solidified mathematical foundations for stochastic theory, based on, among other, previous works by Griffiths and Tavareé [23] as well as a birth-death process. Exponents different from −2 started recently appearing in simulations and empirical observations, such as e.g., in [32], and these were Tung and Durrett [41] who put a special-case model on a solid stochastic process footing.

We proceed as follows. Consider a population of dividing cells with a positive growth rate *g*(*N*), where *N* (*t*) is the population size at time *t*. Suppose that mutation rate (per time) is equal to *h*(*N*). Let *M* (*t*) represent the number of mutations that occurred up to time *t*. Mathematically, this means

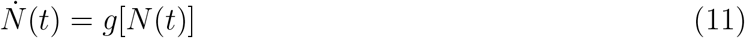

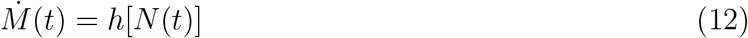

Equation (11) has an implicit solution

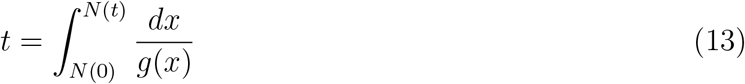

Under neutrality, variant frequency *f* is equal to the reciprocal of the population size at the time *t*_*f*_ when the mutation occurred:

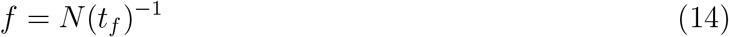

The latter equation has a unique solution if *g*(*N*) *>* 0. The tail *T* (*f*) of the SFS, i.e., the number of variants with frequency greater than *f* is equal to the integral of 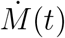 from 0 to *t*_*f*_ :

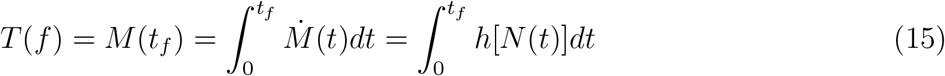

##### Proposition

Let us assume that *g* : ℝ_+_ → ℝ_+_ *\* {0} so that *N* (*t*) increases with *t*. Then the cumulative tail of the SFS has the form

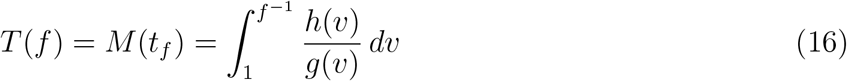

##### Proof

Under the hypothesis, we can substitute *v* = *N* (*t*), yielding 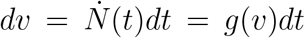. In addition, based on Equ. (14), *N* (*t*_*f*_) transforms into *f* ^−1^, hence the assertion.

##### Corollary

SFS equals *S*(*f*) = − *dT* (*f*)*/df*. We can restate the expression (16) by substituting *φ* = *v*^−1^ under the integral. This will result in

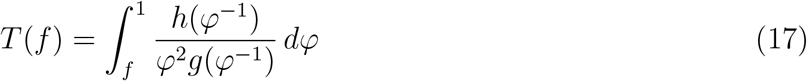

and

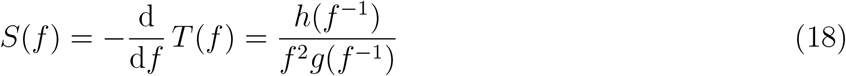

##### Conclusions

1. If *g*(*v*) = *rv*^*β*^, and *h*(*u*) = *µu*, where *r* and *µ* are correspondingly growth and mutation rates,

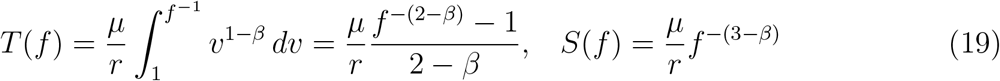 Let us notice that the solution of Equ. (11), with *N* (0) = 1 has the form

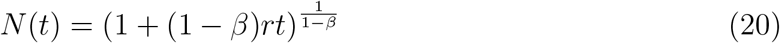

if *β* ∈ (0, ∞) *\* {1}, with *N* (*t*) = exp(*t*) if *β* = 1. In other cases, we may have *N* (*t*) *↑* ∞ as 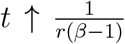, i.e., population explodes in finite time. However, since 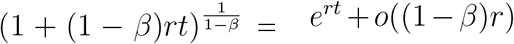, then if |1 − *β*|*r* is small, the expression (21) becomes close to the exponential function, while the explosion time recedes to ∞.
2. Explosion of solution, which is biologically impossible, can be avoided by applying a logistic term in the growth rate function, e.g., by replacing the right-hand side of Equ. (11) by

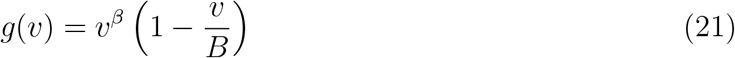

where *B* is the carrying capacity (ultimate limit on tumor size). In this case the explicit solution does not generally exist, except for integer *β*. However, computations prove the logistic term affects the SFS only for *f* -values below certain threshold.
3. In Conclusion 1, SFS has the power tail with exponent −*α* = *beta* − 3, i.e., it is equal to −*α* = 2 if *β* = 1, consistent with the exponential population growth.
4. Based on Conclusion 1, the estimate of *µ/r* can be constructed as

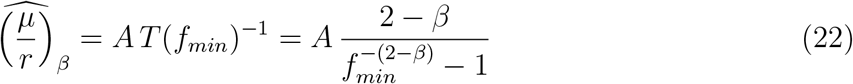

where *f*_*min*_ = 1*/n* is the lowest *f* for which SFS can be theoretically computed with sample size (sequencing depth) *n*, and 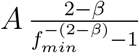 is the “correction for sample size and non-exponentiality” term, which becomes equal to 1*/*(*n* − 1) ≈ 1*/n* if *β* = 1, i.e., identical in practice to the correction in Equ. 10. If we use the model with logistic term, the correction is still approximately valid.

## 3 Results

### 3.1 Comparison of impact of removal of FFPE-induced spurious mutation calls, using different bioinformatics tools

#### 3.1.1 Number of variants per sample before and after filtration

Mutational variants from the BRCA cohort were filtered first by coverage (≥ 10) making such prepared samples “unfiltered” reference point. The average number of variants per sample was close to 7500 in both, primary tumor and lymph node. Comparing to the number of variants seen in TCGA database (on average ca. 450 variants per sample), this number is unusually large. Using bioinformatics tools we attempted to filter out artifacts of formalin fixation. The proportional share of three categories: SNV, artifact and uncategorized among unfiltered variants are shown in Figure 4. After filtration with SOBDetector still a large proportion of variants was retained (Figure 4, upper panel and Fig. S4), with the average of ca. 4000 variants per sample. FFPolish, in turn, left a small number of variants (Figure 4, middle panel and Fig. S4), on average 141 in primary tumor (P1) and 124 in lymph node sample (L1). DEEPOMICS and DEEPOMICS plus retained on average 200-250 variants (Fig. S4). The difference of frequencies of possible categories between tools results from the differences in their algorithms, with SOBDetector also marking part of variants (among others insertions and deletions) as “uncategorized”. See [37] for more details.

**Figure 4:**
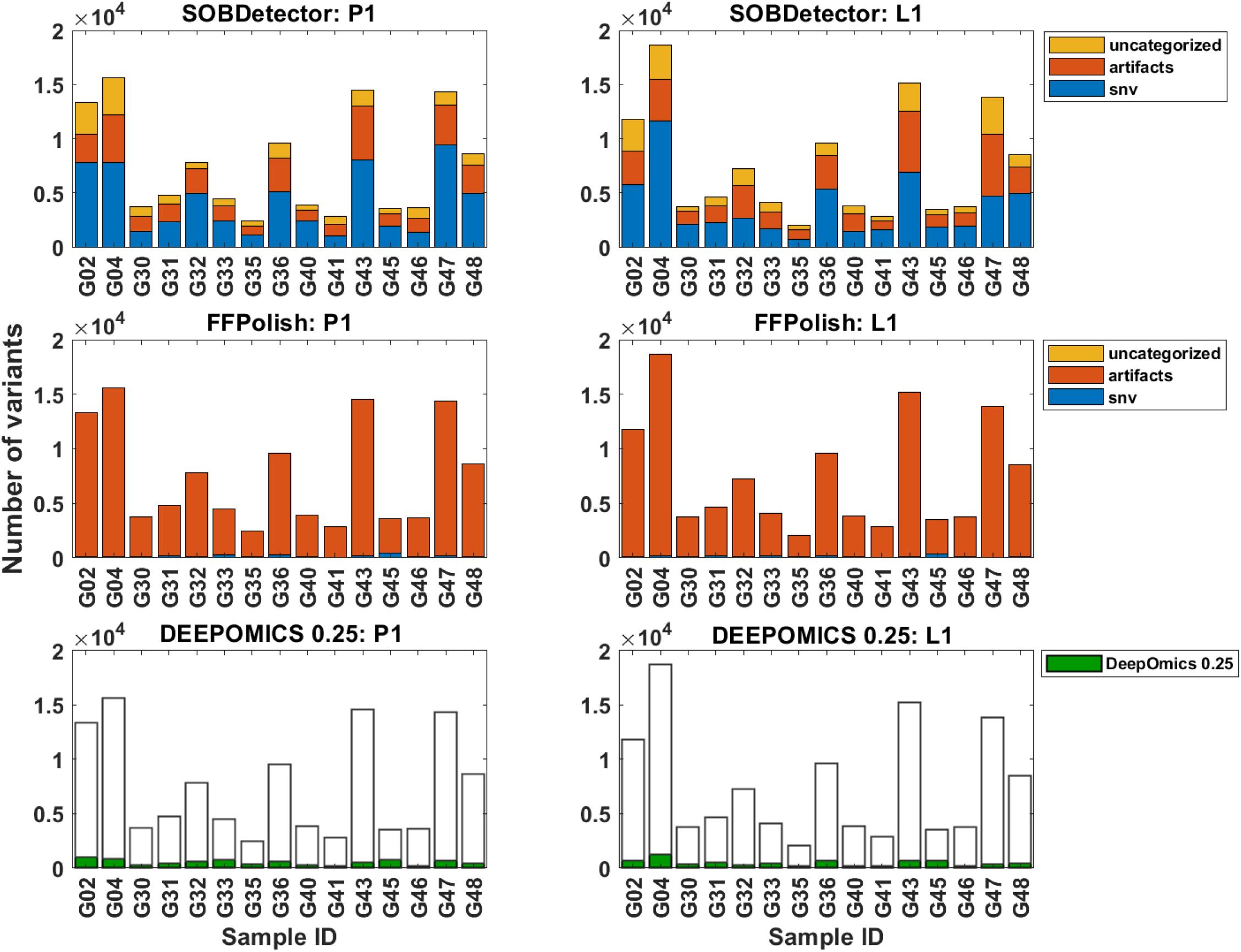
Numbers of all variants per sample with marked categories assigned with three chosen protocols of filtration for primary tumor (P1, left column) and lymph node (L1, right column). Bar charts for SOBDetector (upper panel) and FFPolish (middle panel) represent share of the three categories: snv, artifacts and uncategorized in total count of variants. Bar chart for DEEPOMICS with cutoff FFPE score adjusted to 0.25 shows number of variants retained after filtration with respect to the total count of variants.

#### 3.1.2 Filtration efficacy

To evaluate filtering efficacy we examined the counts of single nucleotide variants (SNV’s) belonging to respective nucleotide substitution categories (Figure 5). Because of the fact that we do not have fresh-frozen counterparts of our samples we compared them to the samples from fresh-frozen breast cancer material from the TCGA database, taking it as an approximate reference point. The results shown in Figure 5 clearly show significant enrichment of or sequencing data with mutations characteristic of formalin fixation as a method of sample preservation. Moreover, these results show that SOBDetector failed to remove most of the artifacts. The results obtained with FFPolish filtration were better and the most effective was the DEEPOMICS software, for which the proportional share of variants resembles that seen in TCGA.

**Figure 5:**
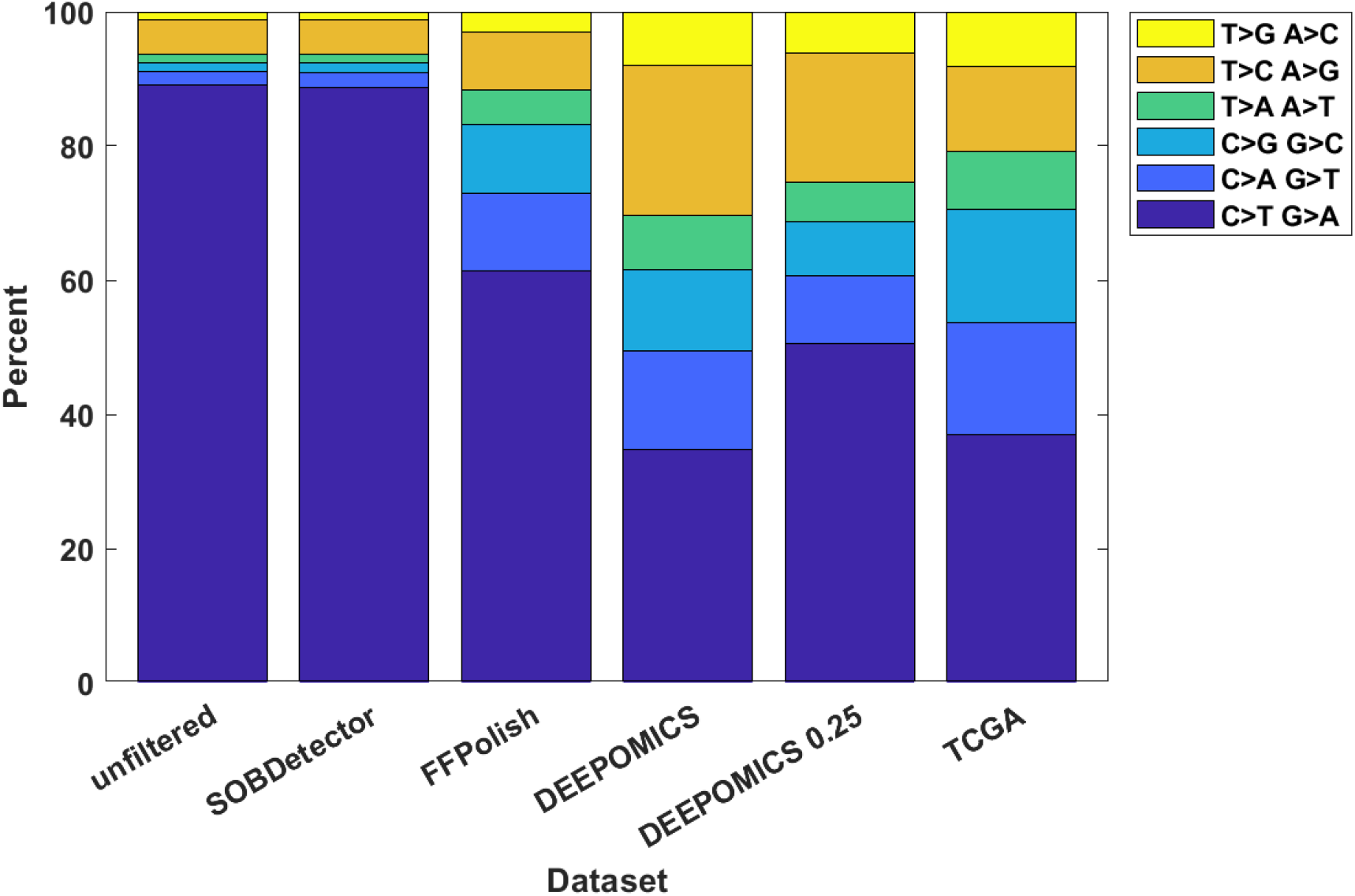
Filtration efficacy of FFPE-removing software in comparison with TCGA breast cancer data. Shown is the percentage of variants belonging to particular categories. Complementary base substitution were merged together for brevity.

#### 3.1.3 DEEPOMICS results after FFPE score threshold adjustments

Due to the fact that DEEPOMICS was the most effective in variants removal, we chose this software for further analyses. However, the overall low number of variants after filtration made it difficult to perform some of them. Therefore, we adjusted the default threshold value of FFPE score (0.5) to lower value (0.25). Results of the adjustment are shown in Figure 4 (bottom panel): the average number of variants per sample increased to 453 in primary tumor and 511 in lymph node samples. Filtration efficacy decreased, but it was still far better than the outcome of the other tools (Figure 5).

### 3.2 Analysis of frequencies of driver mutations

We tallied the mutational hits across the whole cohort in genomic regions encoding known breast cancer driver genes (Figure 6). Analyzed were only mutations which impact protein structure (missense, frameshift etc. variants) and have variant allele frequency (VAF) ≥ 0.05. The most frequently mutated genes differ in cases before and after filtration but are similar in primary tumor and lymph node sample groups. Among the most often mutated driver genes are TP53, PIK3CA and GATA3 in which the mutations belong to the most prevalent in specific subtypes of breast cancer [35, 36]. The overall number of mutations in driver genes was significantly reduced after filtration (from 76 and 81 to 18 and 20 in P1 and L1, respectively), but the extent of reduction (76%) is significantly lower than among remaining variants (93%).

**Figure 6:**
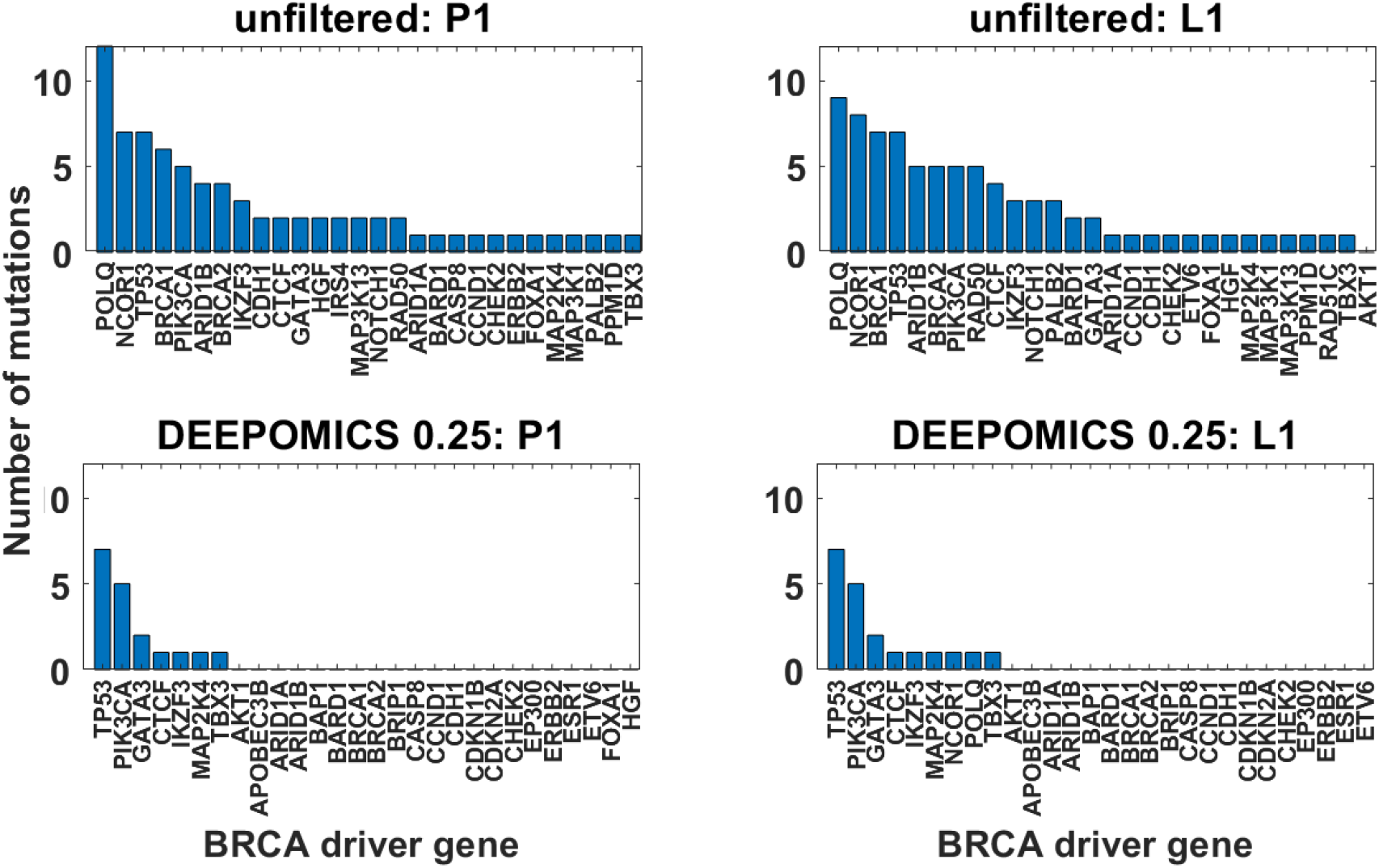
Number of mutational hits across the whole cohort in genomic regions encoding known breast cancer driver genes. Shown are only mutations which impact protein structure and have VAF ≥ 0.05 before (top row) and after the filtration with DEEPOMICS with FFPE score threshold adjusted to 0.25 (bottom row) for primary tumor (P1, left column) and lymph node metastasis (L1, right column).

Next, we determined in which samples the mutational hits occur and how many mutations per sample we observe (Figure 7). The heatmaps show comparison of driver mutation counts before and after filtration with DEEPOMICS with FFPE score threshold adjusted to 0.25. After filtration the most often mutated genomic region of POLQ disappeared and we see only single mutation per driver encoding region per sample.

**Figure 7:**
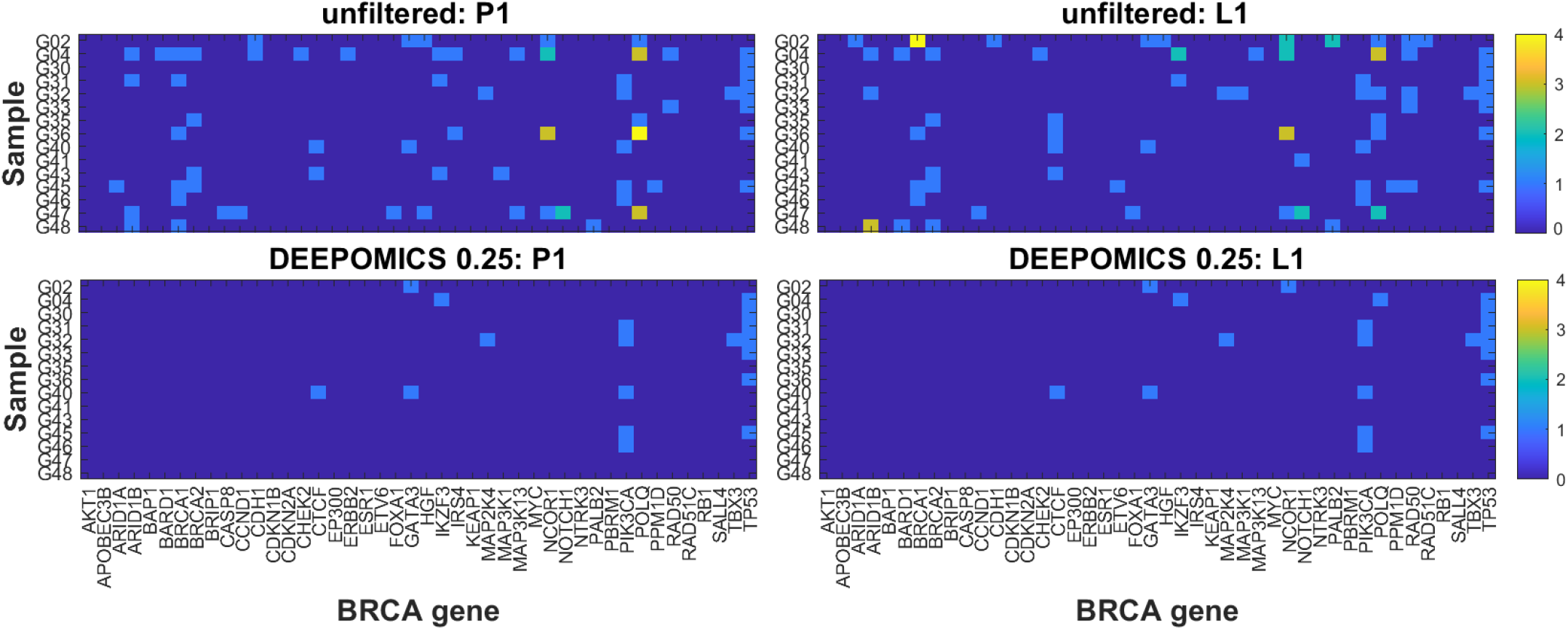
Count of breast cancer drivers per sample before (top row) and after filtration with DEEPOMICS with FFPE score threshold adjusted to 0.25 (bottom row). Shown are results for primary tumor (P1, left column) and lymph node metastasis (L1, right column). Color corresponds to the number of mutational hits in genomic region encoding particular driver gene.

Finally, we compared how many mutational hits are shared by both primary and lymph node (Figure 8). Before filtration, some mutations were observed in both of the paired samples but there was also a number of mutations which were unique to the primary tumor or lymph node metastasis. Filtration with DEEPOMICS left intact mostly mutations present in both P1 and L1, and only two mutations unique to lymph node metastasis.

**Figure 8:**
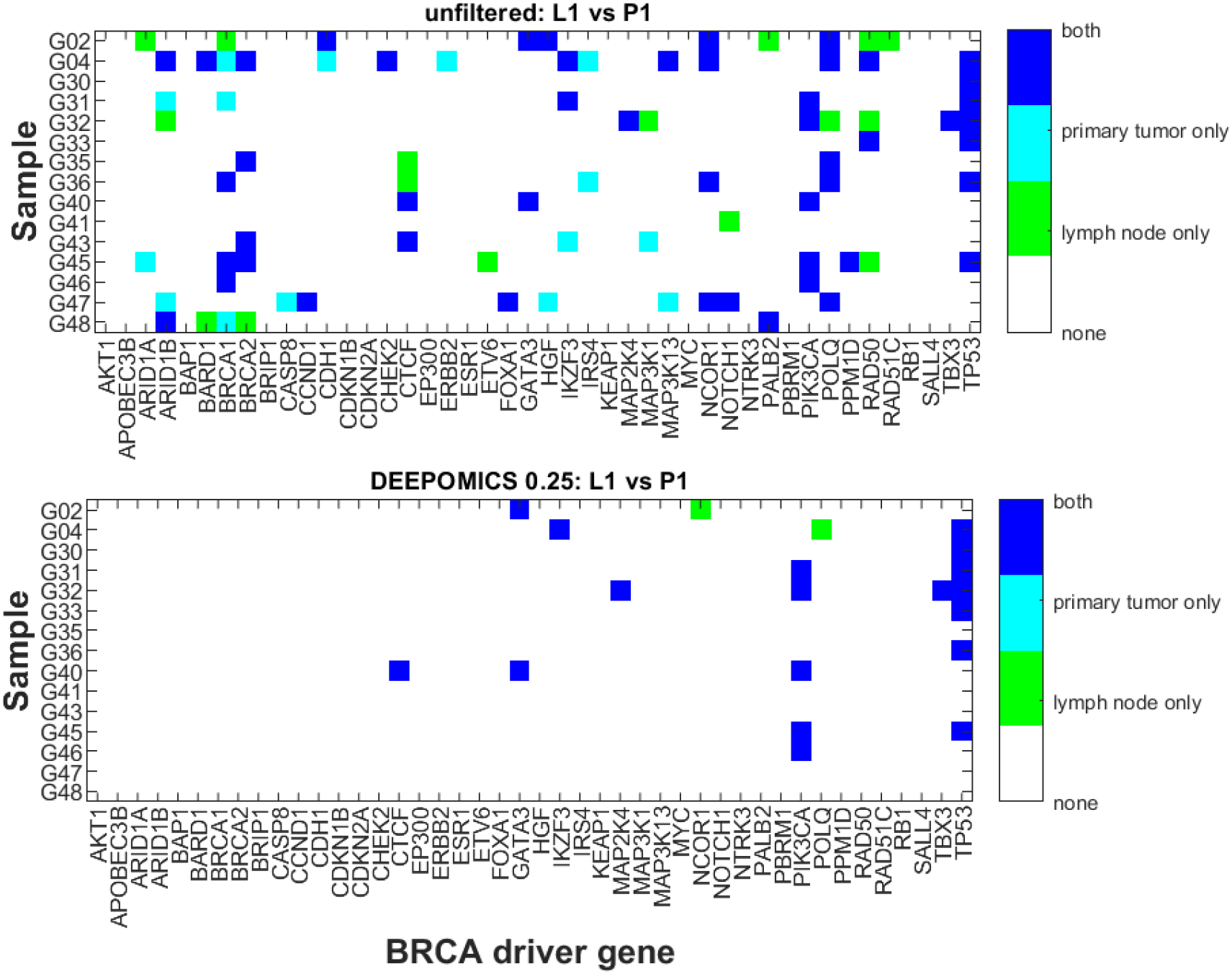
Driver mutations present in primary tumor or lymph node across samples before and after filtration with DEEPOMICS with FFPE score threshold adjusted to 0.25. The color corresponds to the category: overlapping mutations (dark blue), mutations present only in P1 (cyan) and mutations present only in L1 (green).

In the data filtered with DEEPOMICS TP53 mutation was present in all BRCA subtypes (except luminal A), PIK3CA was present in all subtypes with the smallest count in TNBC, we observed also two mutational hits in GATA3 and single hits in CTCF, MAP2K4, NCOR1 and TBX3, all consistent with the results presented in [36] and [34]. Additionally we observed one mutation in IKZF3 (found also as often occuring in HER2+ cancers [31], as in our cohort) and in POLQ associated with poor prognoses in breast cancer patients [29].

### 3.3 Oncogenes and tumor supressors at variable CNV Levels

We carried out joint estimation of copy number variation (CNV) and fraction of tumor cells (purity, *ρ*) in all 31 specimens, using the ASCAT software. As it is known, ASCAT, and other equivalent methods can provide two or more nearly equivalent estimate sets. The reason is that a cell population which is a mixture of normal cells (diploid) and tumor cells (variable CNV) may be barely distinguishable based on DNA sequencing from another one that has a lower purity but a higher CNV. In such doubtful cases, we decide by coordinating of average CNV levels of the P1 (primary) and the L1 (invaded lymph node).

Figure 9 depicts ASCAT estimates of DNA CNV (ploidy) levels in segments of chromosomal regions, for all specimens analysed in the study. Certain patterns can be found, such as the predominant gains in specimens G02, G04, G30, and G31 (HER2+ tumors), losses in specimens G32, G33, and G35 (TNBC tumors), and mainly diploid genomes in G40, G41, G43 and G46 specimens (luminal A tumors). Due to limitations of the size of our sample these findings are not systematic, but low copy number alterations were reported for luminal A subtype [36] while luminal B were found characterized by complex landscape of copy number changes [34].

**Figure 9:**
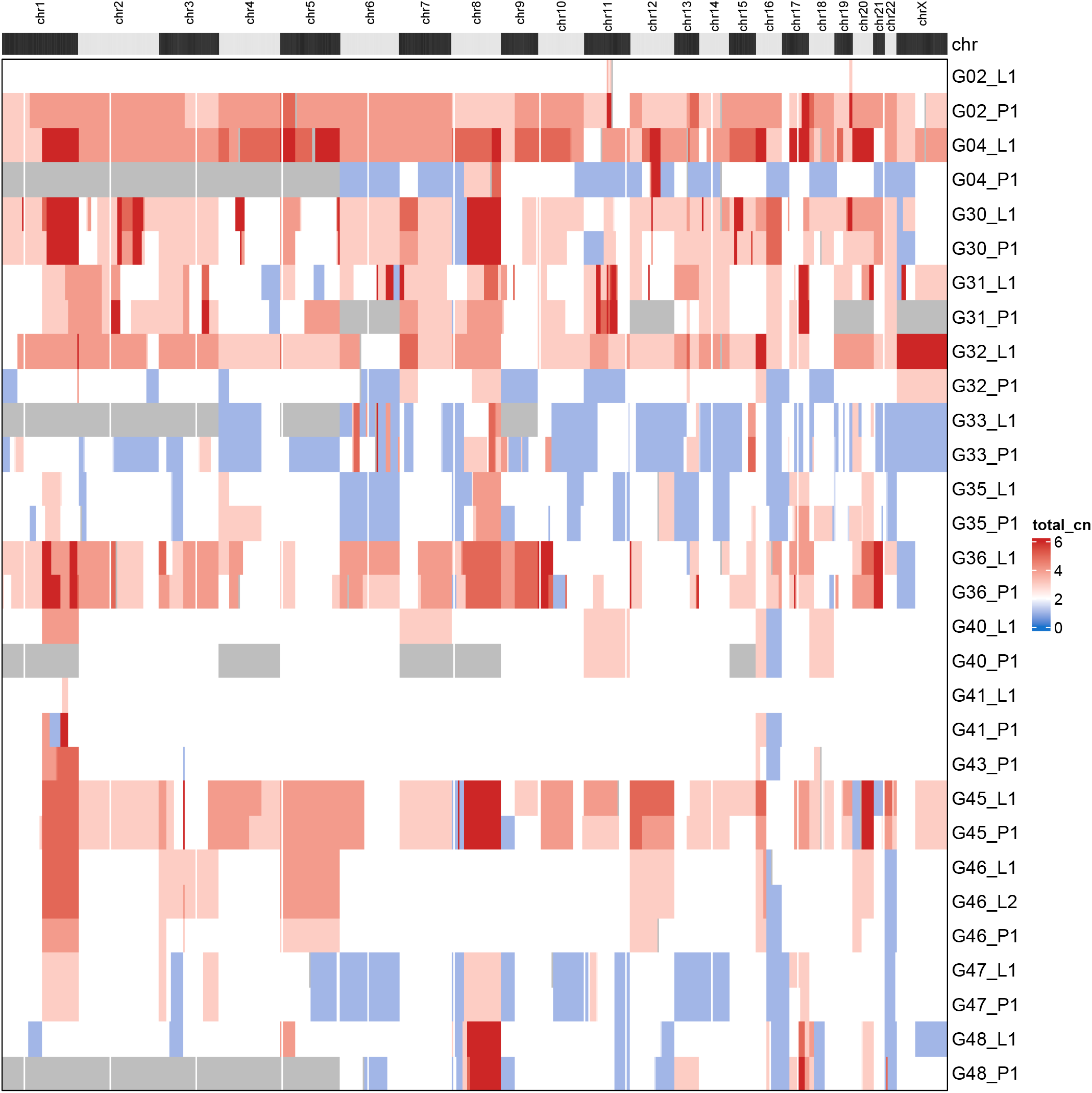
ASCAT estimated DNA CNV (ploidy) levels in segments of chromosomal regions, for all specimens analysed in the study. Red color corresponds to copy number gains and blue color to copy number losses. Please notice the predominant gains in specimens G02, G04, G30, and G31 (HER2+ tumors), losses in specimens G32, G33, and G35 (TNBC tumors), and mainly diploid genomes in G40, G41, G43 and G46 specimens (luminal A tumors).

The main reason for CNV analysis at the genomic scale is to understand the pattern of genome evolution. For example, if a given driver is present at the region of CNV twice the normal (i.e., ca. tetraploid, or 4), but its VAF equals 0.25, then this means that the corresponding nucleotide substitution occurred after this portion of cell’s genome was duplicated. If the VAF equals 0.75, this may mean that nucleotide substitution before duplication, but then one copy of the region in question was lost, etc. Dissecting relationships such as this may facilitate determining the order in which the CNV of different parts of the genome evolved.

Another aspect of the problem is selection for the oncogenes (OG) and tumor suppressor genes (TSG). Assuming that a mutated OG positively affects cell proliferation, this action is increased if it exists in an increased copy count. The same, but to a different extent may be true for a mutated TSG. Therefore by the action of selection, the ratio of mutated OG to TSG elevated or reduced in the regions of increased CNV, with the opposite hypothetical trend being present in the regions of reduced CNV. A thorough analysis, based on a number of cancer types is summarized in Figure 2 in reference [43]. These authors show empirically the positive correlation of CNV with elevated OG to TSG ratio.

We carried out a slightly different, but equivalent analysis, based on BRCA COSMIC cancer genes [40], relating the CNV of the region harboring given drivers to either the regular ploidy (= 2) or to the average ploidy of a given tumor. The analyses were carried out in two versions: Using unfiltered calls which have a strong admixture of FFPE-related artifacts, and using calls filtered by DEEPOMICS software [26], which seems to correctly restore the mutational spectrum in BRCA to that typical of frozen non-FFPE specimens from TCGA.

Details are depicted in a series of figures in this section and in the Supplement. Figure 10 shows the result of analysis of copy number in genomic regions containing driver genes in all samples before (top panel) and after filtration with DEEPOMICS with FFPE score threshold adjusted to 0.25 (bottom panel). The background dashes (horizontal for P1 and vertical for L1) correspond to the relative copy number in the region with respect to assumed normal ploidy, where “down” (blue) represents copy number loss vs “normal” (green) ploidy and “up” (red) represents copy number gain. The assumption of “normal” genome ploidy equal to 2 is not very realistic in the case of BRCA cancer for which frequent copy number gain and very heterogeneous copy number profiles were reported [35, 34]. We calculated the average copy number in three major groups of genomic regions: containing tumor suppressor genes (TSG), oncogenes (OG) and OG/TSGs. Our results (Figure 11, top panel) indicate that the average copy number in all three groups is higher than 2 (with the highest values in the oncogene group). Therefore, in parallel, we conducted an analysis with baseline ploidy of the sample estimated as weighted average across genomic regions in this sample (see Figure S5). The plot area of both figures is divided into three major groups: genomic regions for tumor suppressor genes, oncogenes and a cluster of genes acting in different ways depending on specific context, such as e.g., a BRCA subtype or a specific mutation type. In both figures a noticeable increase of copy number (prevalence of red marks) is seen in regions containing oncogenes. Assuming baseline ploidy calculated for specific samples, which is on average equal to 2.4 in primary tumor and 2.6 in lymph node metastasis, this effect is not that prominent (due to a general increase in ploidy). In this case, in turn, the decrease of copy number in regions encoding tumor suppressor genes might be noticed (Figure S5). Also the region containing the TP53 gene (here classified as a mixed-role gene) is characterized by a lower than baseline copy number; this effect is shared among most of the samples which additionally have mutational hit in this gene. These results are consistent with literature reports which show also amplifications in regions containing PIK3CA, FOXA and ERBB2 (encoding HER2 protein) genes and copy number loss in regions containing RB and MAP2K4 [4].

**Figure 10:**
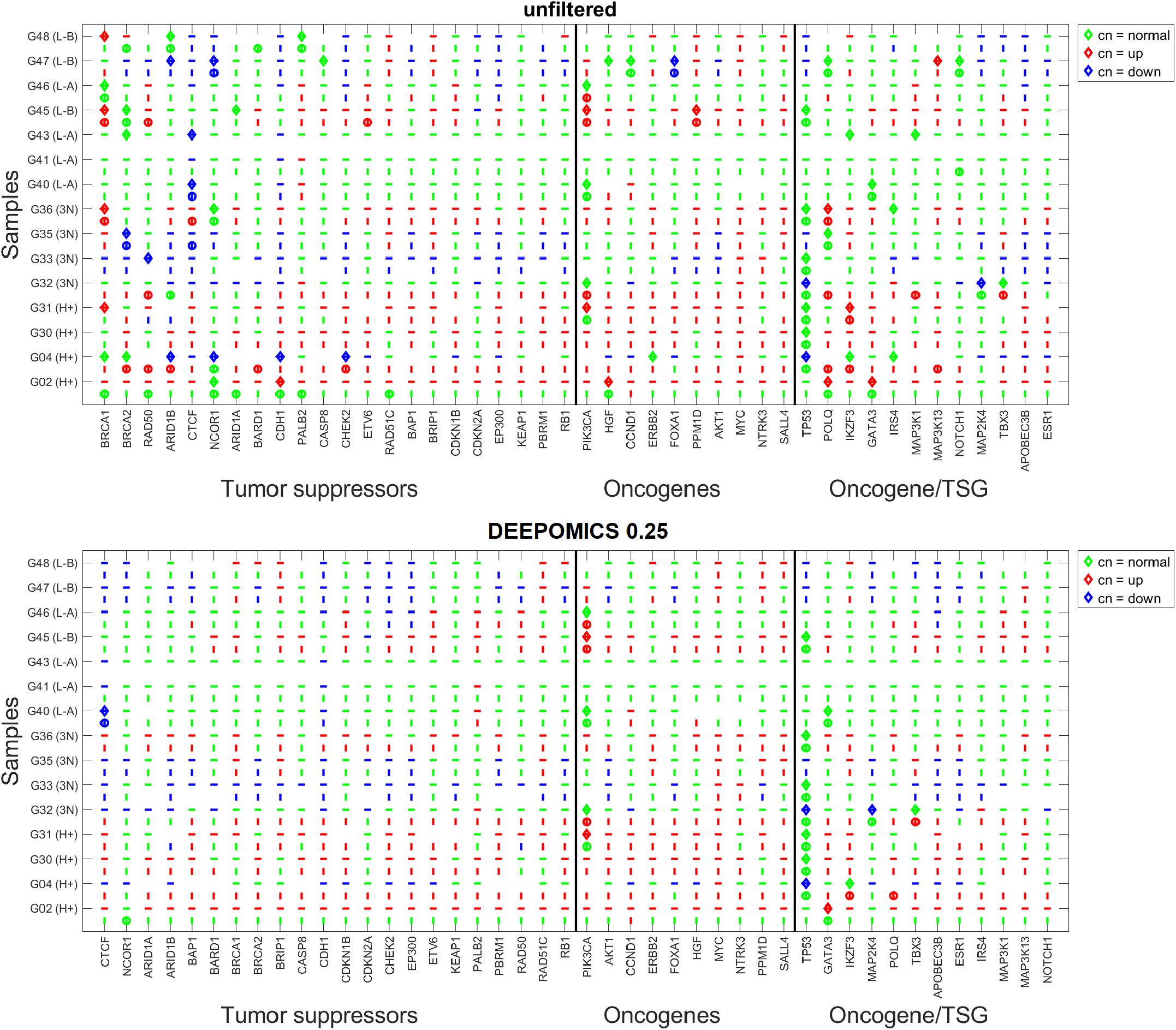
Ploidy in genomic regions encoding driver mutations in all P1 and L1 samples before (top panel) and after filtration with DEEPOMICS with adjusted FFPE score threshold (bottom panel). Dashes (horizontal for P1 and vertical for L1) represent relative ploidy in region with respect to assumed normal ploidy equal to 2. The change in ploidy is encoded with color: green - normal ploidy, red - increased ploidy, blue - decreased ploidy. Additionally mutations in driver genes are marked by diamond (P1) and circle (L1). Horizontal axis of the graph is sorted by driver gene function and divided into 3 major groups: TSG’s, OG’s and OG/TSG’s. In each of the major groups driver genes are sorted in descending number of mutational hits.

**Figure 11:**
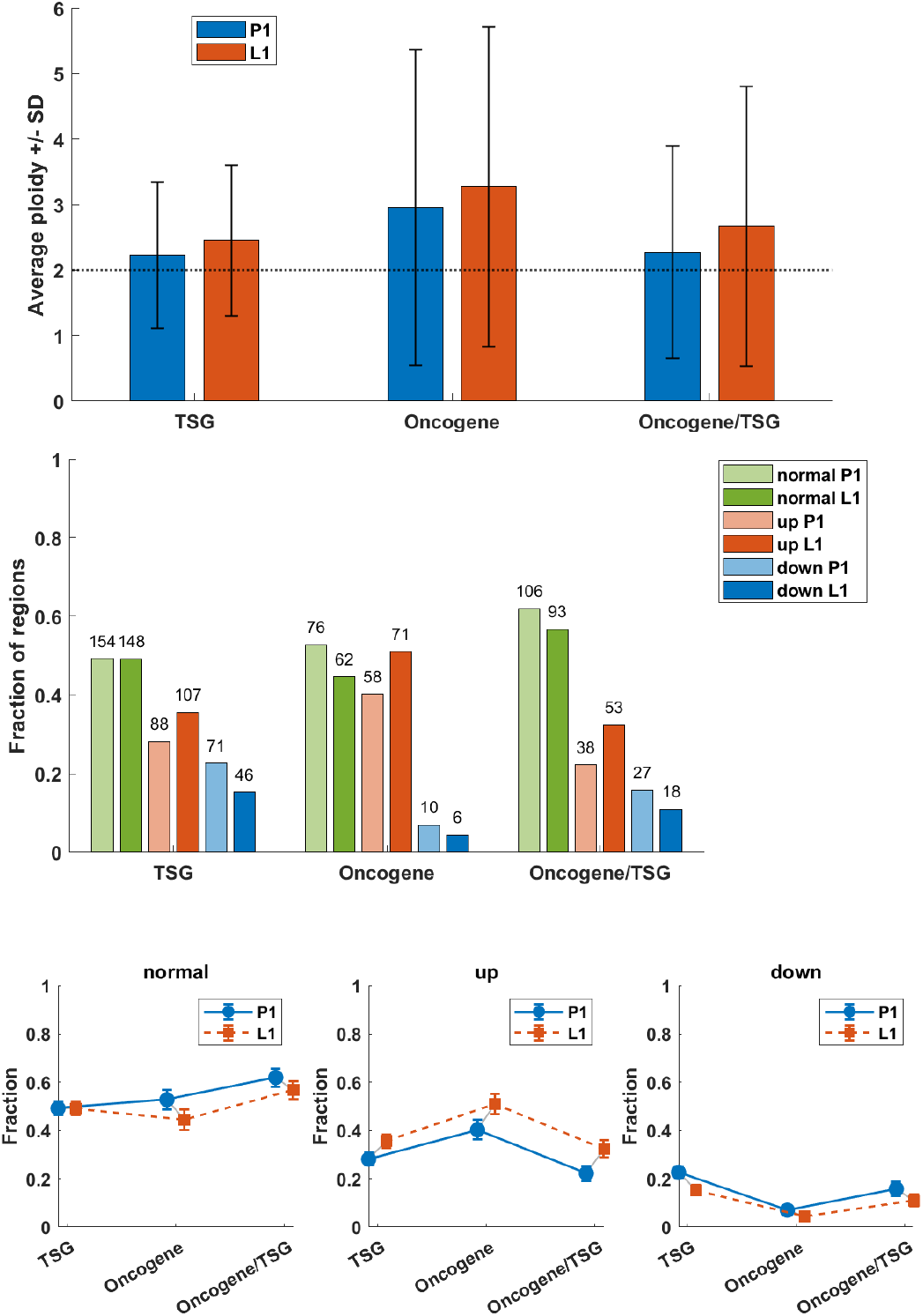
Statistics of copy number in regions encoding driver mutations across all P1 and L1 samples, assuming normal ploidy equal to 2. **Top panel:** Average copy number in regions encoding three major groups of driver genes: TSG, OG and OG/TSG. **Middle panel:** Fractions of regions with copy number increased (red), decreased (blue) or unchanged (green) with respect to normal ploidy assumed equal to 2. The height of the bar depicts proportional share in given major group, while the numbers above are the counts of regions of a given type. **Bottom panel:** Fractions of regions encoding driver mutations in P1 and L1 with copy number unchanged (normal), increased (up) and decreased (down) in the three major groups of driver genes. Distribution of TSG and oncogenes OG in P1, comparing copy number down vs. normal vs. up (counts: down — TSG 71, OG 10; normal — TSG 154, OG 76; up — TSG 88, OG 58): *χ*^2^ = 18.6, *p <* 0.001. In L1 (counts: down — TSG 46, OG 6; normal — TSG 148, OG 62; up — TSG 107, OG 71): *χ*^2^ = 15.8, *p <* 0.001. Both results are significant — TSGs are enriched among copy loss and OGs are enriched among amplifications. Total counts of regions with copy number down/normal/up (TSG, OG, and OG/TSG combined) across P1 and L1 (counts: down — P1 108, L1 70; normal — P1 336, L1 303; up — P1 184, L1 231): *χ*^2^ = 14.7, *p <* 0.001. The result is significant — P1 shows relatively more copy losses while L1 shows relatively more amplifications.

We determined how many BRCA-driver containing genomic regions fall in the category of increased, decreased and normal (unchanged) copy numbers. Results are subdivided into three major groups: TSG, OG and dual effect genes OG/TSG. In all three groups, the fraction of regions with increased copy number is higher than that with decreased. In the TSG group half of the genomic regions have copy number equal to 2 (Figure 11, middle and upper panel); this value is similar in OG group and slightly higher in OG/TSG cluster. The percent of regions with increased copy number is similar between TSG and OG/TSG groups, while in OG group it is increased. The fraction of regions with decreased copy number is the highest in TSG, moderate in OG/TSG and the lowest in OG groups. For each sample we tested independence between copy-number category (down vs. normal vs. up) and other groupings using the chi-square test for 2 *×* 3 contingency tables. Reported values are the chi-square statistic (*χ*^2^) and two-tailed *p*-value. Distributions of number of regions containing TSG and OG in P1 and L1, with copy number decreased, normal and increased have shown that in both groups regions containing OG are significantly enriched among amplifications, while regions containing TSG are more frequent among (loss of copy).

This trend, which is most clearly shown by Figure 11, remains present also in the case of baseline ploidy of the samples estimated as the weighted average of genomic regions in these samples (Figure S6, bottom panel) but the fraction of regions falling to each category shifts towards unchanged (normal) and decreased (down) diploid in this region. Except for the OG group, there are more regions with copy number lower than baseline than the opposite.

There exists a significant difference in the distribution of copy number in regions encoding driver genes between primary tumor and lymph node samples. In general, lymph node samples have higher average copy number, overall and in regions containing driver genes.

### 3.4 Variant allele frequency (VAF) and cancer cell fraction (CCF) spectra

As mentioned in Methods, we carried out analyses of two types of spectra: SFS and CCF, in both cases using data either unfiltered and hence including FFPE-related artifacts, or filtered using DEEPOMICS software [26]. We present a review in the next subsection.

In addition, we carried out evolutionary analysis using a novel tool developed by Khanh N. Dinh, based on a probabilistic methodology in [11] (see Methods for detail). Results and conclusions of that analysis are presented in a further subsection.

#### 3.4.1 Review of the unfiltered and filtered spectra

Since the mutation pattern of our samples is highly heterogeneous, we decided to present the spectra of individual samples in the Supplement. In this section, we present aggregates of all samples. Please note that in the model such as that outlined in the Methods, a typical spectrum of mutations in the expanding cell population should have two or more overlapping components: The first counting from the left, corresponds to the neutral mutations from all cell clones, while the centrally located “hump” (at frequency close to 0.5 after correction for purity) corresponds to the truncal mutations inherited from the ancestral cell of the tumor. Further humps may correspond to additional subclones. The neutral component is theoretically a decaying power curve, but it is frequently corrupted by removal of the lowest-frequency data as a result of data-cleaning [11].

In the aggregate spectra as depicted in Figure 12, we distinguish a very strong “neutral” component in the unfiltered data, but mostly the dominating truncal and subclonal humps in the filtered ones (panels filtered using the DEEPOMICS software under different settings). The most interesting shape of all graphs exhibits the spectrum generated using the FFPolish software. However, the overall low number of post-filtration variants and moderate efficacy in filtering out the characteristic artifacts of formalin fixation excludes it from further analysis.

**Figure 12:**
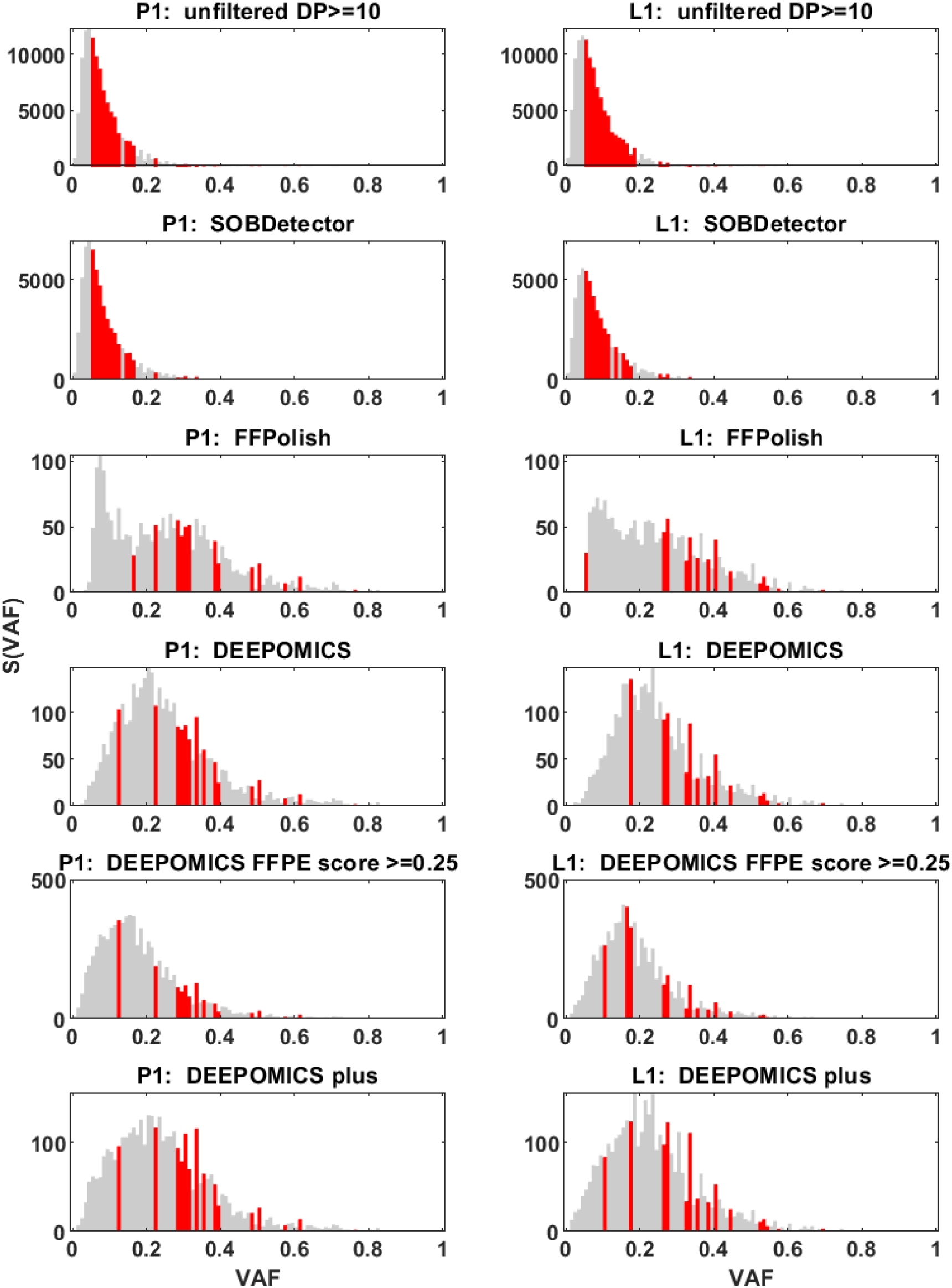
Site frequency spectrum (SFS) based on aggregated variant allele frequency (VAF) from patients samples from primary tumor (P1, left column) and lymph node metastasis (L1, right column). With red marked are bars for VAF thresholds in which mutation variants in driver genes were observed. Each panel from top to bottom was generated using different filtration method. Note the differences on Y axis between methods.

In addition to the shape of aggregated spectrum Figure 12 shows also the position of driver genes in the frequency spectra. Marked with red are bars for VAF frequencies at which mutation variants in driver genes were observed. The count of driver genes in particular VAF thresholds is shown in a corresponding Supplementary Figure S7 in the semi-logarithmic scale.

To take into account purity and ploidy which vary across samples, we calculated Cancer Cell Fraction (see Methods) and presented it in a form of aggregated site frequency spectrum (Figure 13). As in the previous case, the bars present at frequencies where driver mutations were found are marked with red. Corresponding figure in a semi-logarithmic scale showing the number of driver mutations in each bar was included as the Figure S8.

**Figure 13:**
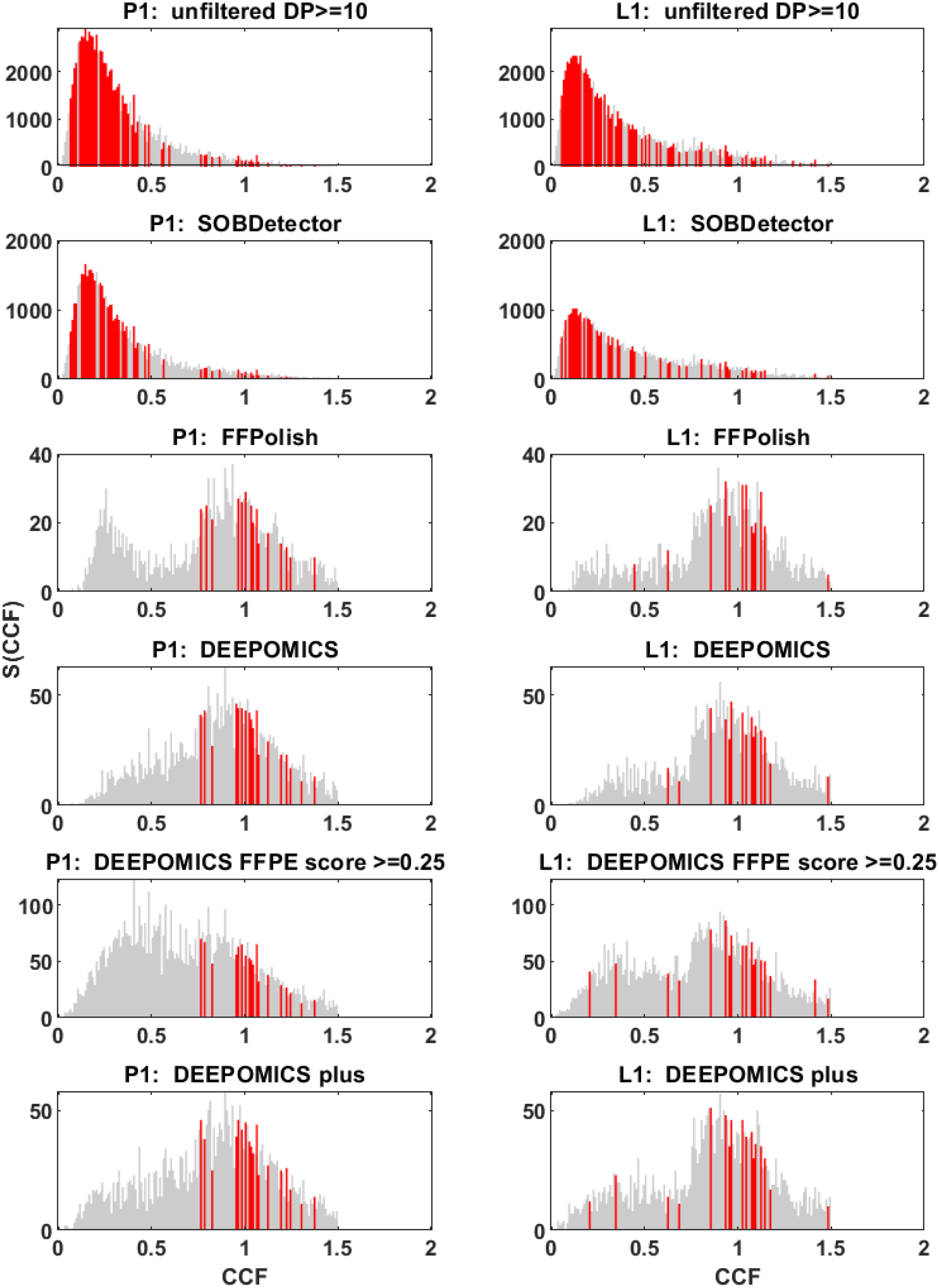
Site frequency spectrum (SFS) based on aggregated cancer cell fraction (CCF) from patients samples from primary tumor (P1, left column) and lymph node metastasis (L1, right column). With red marked are bars for VAF thresholds in which mutation variants in driver genes were observed. Each panel from top to bottom was generated using different filtration method. Note the differences on Y axis between methods and the range of X scale (CCF not rescaled).

In both cases, i.e., in unfiltered and SOBDetector-processed datasets, driver mutations are present in almost all bars with lowest frequency (above 0.05 threshold for VAF imposed on mutations in regions encoding driver genes), while we would expect them to be shared by larger proportion of variant calls. In case of FFPolish- and DEEPOMICS-filtered data, most drivers fall into frequency thresholds from 0.2 to 0.5, which after correction for purity and ploidy and calculation of CCF spectrum shifts them towards the 1/2 of observed frequencies (see cluster of drivers around value 1 in panels depicting FFPolish and DEEPOMICS results).

Another interesting finding concerns the overlaps of mutations between the primary and lymph node subsamples. Results, following DEEPOMICS fitering are presented in Figure 14.

**Figure 14:**
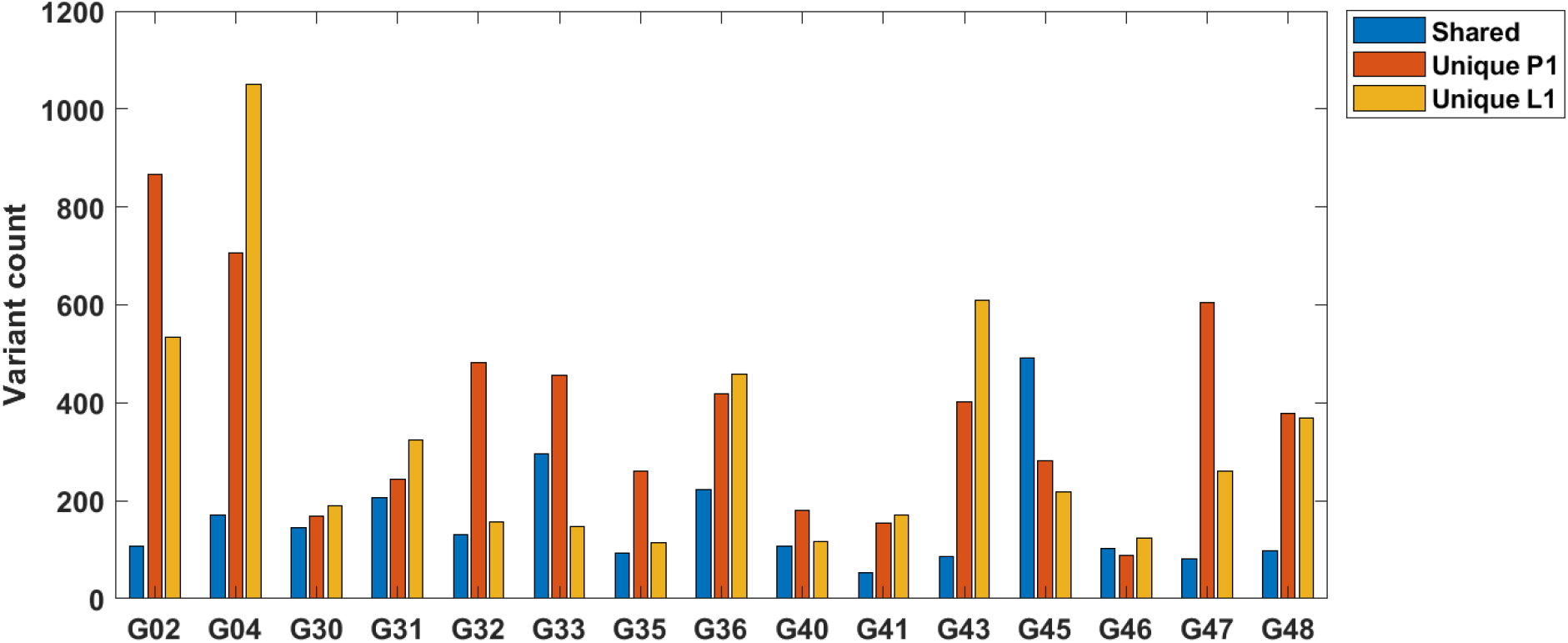
Result of analysis of mutations common in P1 and L1 and mutations unique to these categories in samples following DEEPOMICS filtering. Please note the differences in share of common and unique mutations between patient samples.

#### 3.4.2 Computational analysis of the spectra

We carried out decompositions, as described in Section 2.2.3, of the diploid portions of the SFS spectra of the paired primary (P1) and lymph node (L1) specimens. Because of small sample size partly due to removal of FFPE artifacts, not all of these are informative. However, we identified 3 pairs of specimens, G33, G40, and G41, in which the decomposition led to interpretable results. In Figure 15, the raw SFS from data is depicted by slim black bars, while the grey and brown outlines are the estimated profiles of the neutral power-curve component, and the truncal hump). However, coordinates of the hump and its size, relative to the neutral component vary.

**Figure 15:**
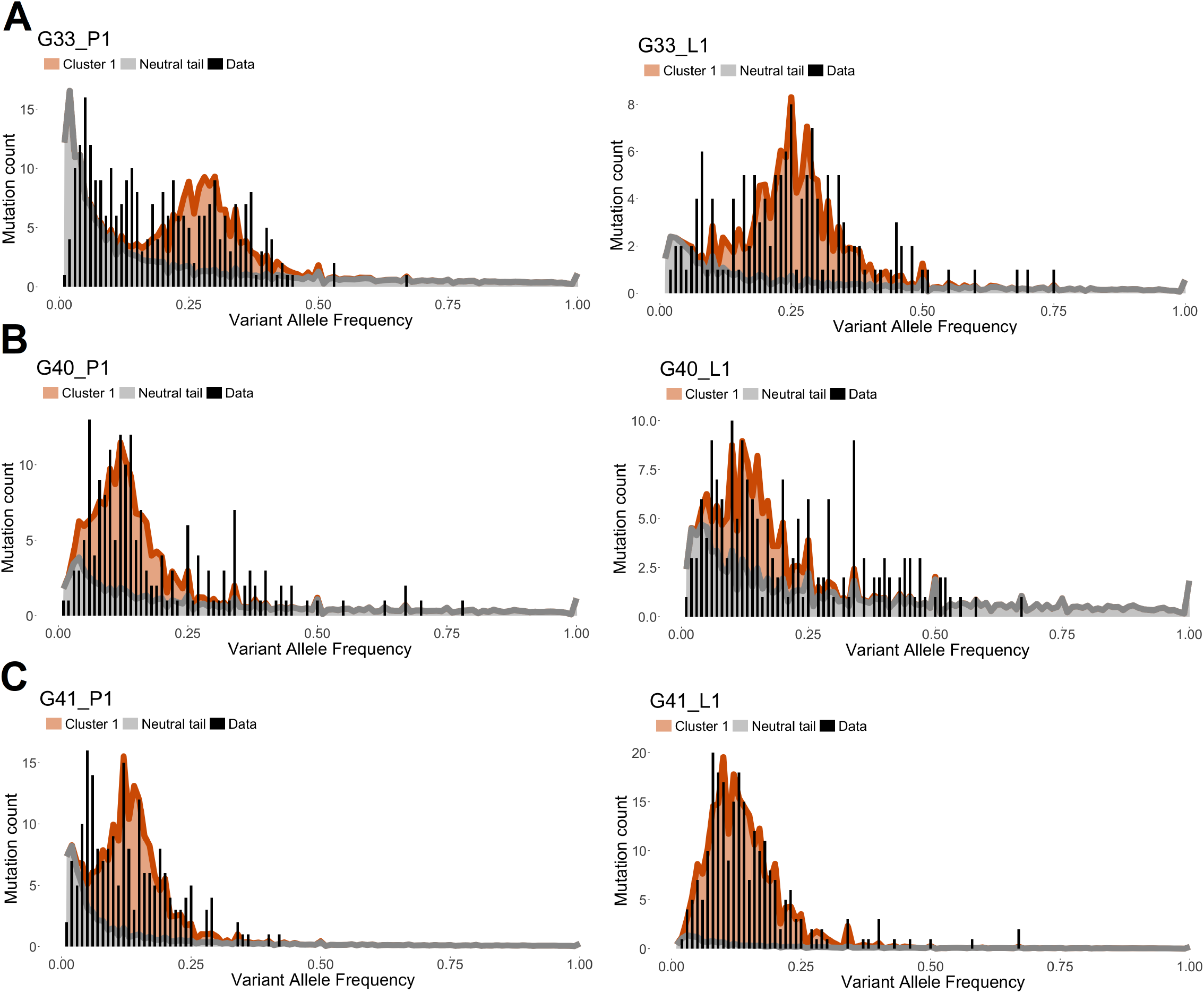
Relationships between the diploid portions of the SFS spectra of the paired primary (P1, *left*) and lymph node (L1 *right*) specimens. We consider 3 pairs of specimens (A) G33, (B) G40, and (C) G41. In each case, the raw SFS from data is depicted by slim black bars, while the grey and brick outlines are the estimated profiles of the neutral power-curve component, and the truncal hump, respectively). However, coordinates of the hump and its size relative to the neutral component vary. For more detail, see discussion in the text.

Estimates of the parameters of the spectrum and some characteristics of the samples are listed in Table 3. Let us note that according to the usual interpretation [44], for regularly diploid genomes the centroid of the truncal hump should correspond to *f*_*h*_ = 0.5 *× ρ* frequency, where *ρ* is sample purity. In other words, the *f*_*h*_*/ρ* = 0.5. However, although for G33 this index is equal to 0.47 (P1) or 0.41 (L1), which is close to 0.5, for G40 it is equal to 0.16 (P1) or 0.15 (L1), and for G41 it is equal to 0.34 (P1) and to 0.13 (L1). We return to this discrepancy in the Discussion,

**Table 3:**
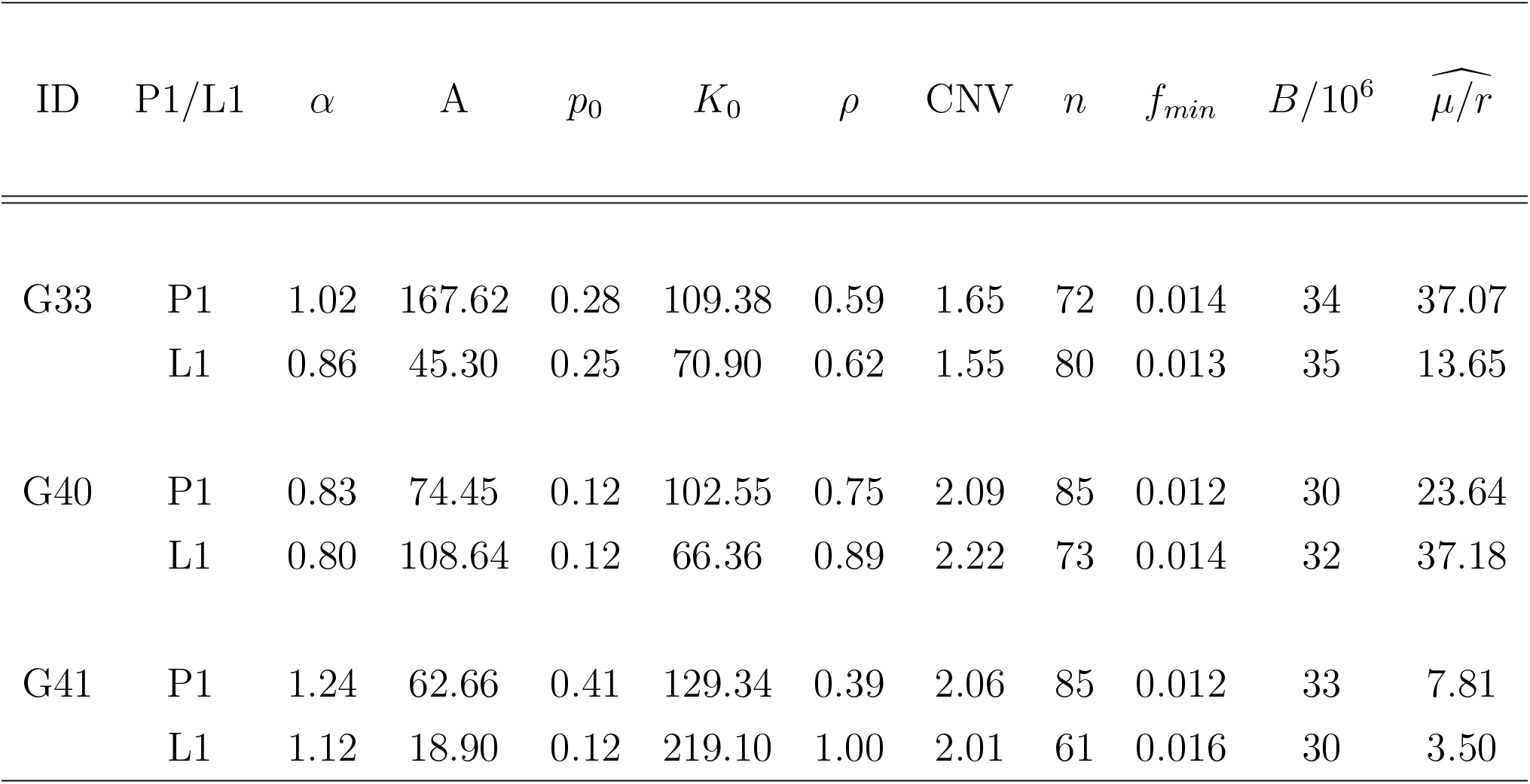
Estimates of parameters for the P1/L1 pairs from patients G33, G40, and G41, based on SFS spectra depicted in Figure 15. Notation: P1, primary; L1, lymph node; *α*, estimated power exponent of the neutral power curve; *A*, estimated neutral mutation count; *p*_0_, estimated centroid of the SFS “hump”; *K*_0_, estimated “hump” mutation count; *ρ*, ASCAT-estimated sample purity; CNV, ASCAT-estimate; *n*, DNA sequencing depth, *f*_*min*_ = 1*/n*, minimum estimable mutation frequency based on the sample, *B/*10^6^, sequenced specimen’s estimated volume in cubic milimeters assumed corresponding to *B* million tumor cells; 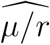, estimated reduced mutation rate.

We focus mainly the on analysis of the identified “neutral” power curve components depicted by black lines in Figure 15. In all cases, the exponents of the power curves vary around −1, in a strong departure from the −2 value, characteristic of the exponential growth. Based on Conclusion 4, Section 2.2.4 this suggests that in early phases of growth, the population of cells with neutral mutations, has been increasing its growth rate as *rN*^*β*^, with *β* = 3 − *α* = 2, which means faster than exponentially. The corresponding estimates of the reduced mutation rate 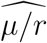, vary from 3.5 to 37.18 mutations per division, which are ca. 2 orders of magnitude higher than in normal exomes.

### 3.5 Mutational signatures

As described in Methods, we performed the analysis of mutational signatures to identify mutational processes likely impacting the tumor samples and also as another layer of FFPE-removal efficacy control (Figure 16).

**Figure 16:**
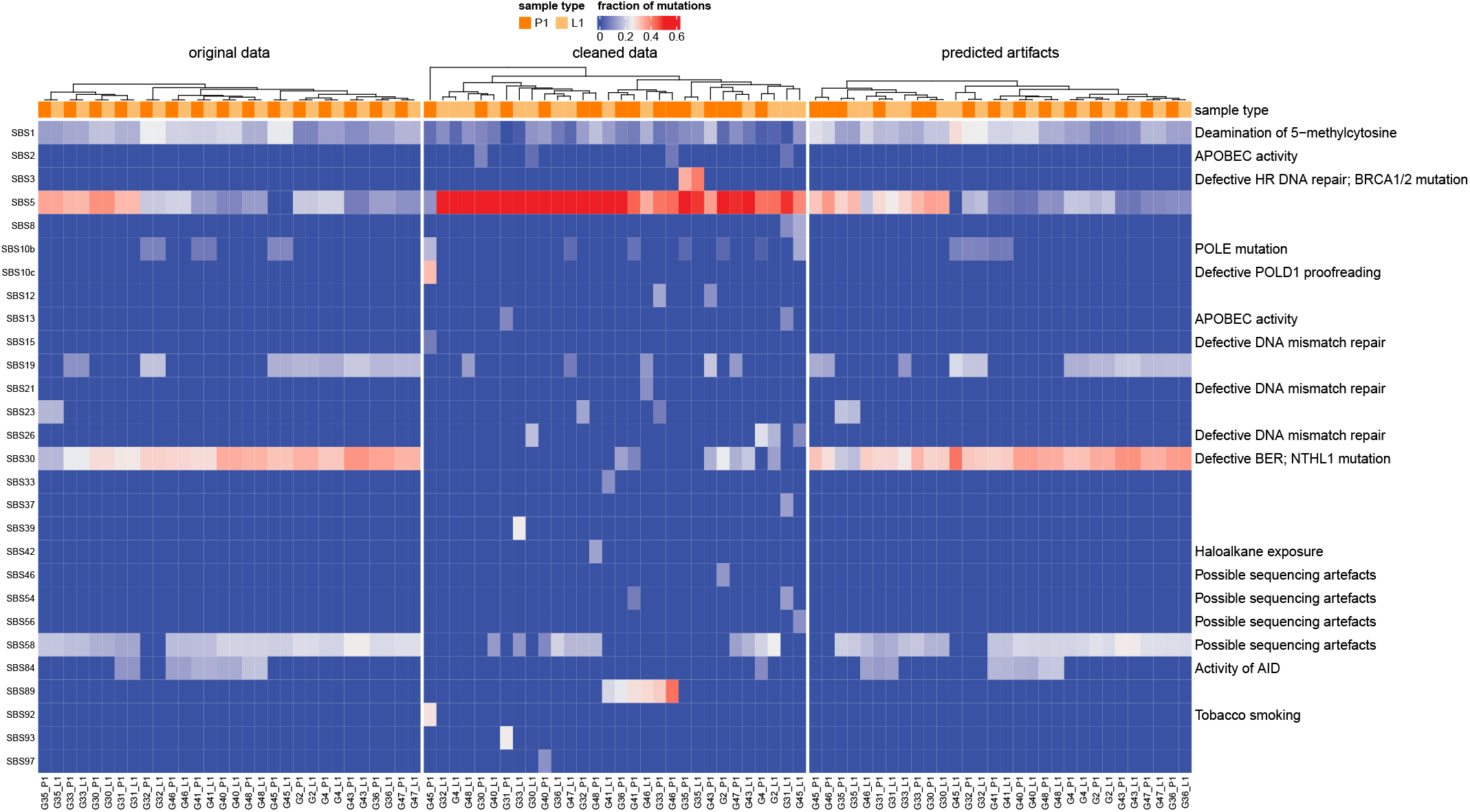
Heatmap of mutational-signature exposures per sample (columns) assigned by SigProfiler. Three panels show unfiltered data, data after filtration with DEEPOMICS (central panel), and in rejected artifacts. Color represents fraction of mutations contributed by each SBS signature.

After filtration, the contribution of signatures identified as “Possible sequencing artefacts” (SBS58) and also SBS30 and SBS1 (the signatures reported to be enriched in FFPE samples [24, 26]) was markedly reduced or eliminated in most samples, while persisted in “predicted artifacts” panel. On the other hand, signatures consistent with biological processes (“clock-like” signatures: e.g. SBS5) remained largely unchanged. This shift increased the relative share of true mutations. A subset of samples, however, still show residual artifact-associated signal after “cleaning”, indicating incomplete removal or low purity of the samples.

## 4 Discussion

In this paper, we carried out analysis of whole-exome DNA sequencing data from FFPE breast tumor specimens collected for diagnostic evaluation of post-surgery follow-up. As is known, FFPE technology of tissue preservation induces DNA alterations that lead to spurious variant calls in sequencing. Therefore, the initial pre-analysis step entails possible identification and removal of the spurious calls. We were unable to use any of the chemical reversal tools currently on the market, but we had at our disposal bioinformatics tools such as SOBDetector [14], FFPolish [16], DEEPOMICS [26] and other, with varying sensitivities and specificities. As discussed in Section 2.1.3, SOBDetector is identifying too few and FFPolish too many variants to match the characteristics of TCGA fresh-frozen tumors. As futher explained in Section 2.1.3 and the forthcoming publication [37], we decided to employ DEEPOMICS with the FFPE score adjusted to 0.25 from the standard value 0.5, to increase the proportion of low-frequency variants, needed to visualize the descending neutral tail.

Based on the results of the analyses of the SNV type distribution (Section 3.1.2) and the mutational signatures (Section: 2.1.5), the method chosen (DEEPOMICS with 0.25 FFPE index threshold) seems to be a good compromise between enrichment in low-VAF variants and the efficacy of filtration.

Our CNV analyses confirm the rough outline of copy number alterations in breast cancer. Purity, ploidy, log-ratio and total copy numbers were identified simultaneously by ASCAT [42]. Average ploidy in the tumors varies in the interval [1.5, 4.52] and with slight exceptions is close for the (P1, L1) pairs. The pairs with highest average ploidy are likely to have undergone whole-genome duplication (WGD) followed by successive ploidy-reducing divisions [13]. Chromosomal landscape of gains and losses is similar as in the cases in a highly cited triple negative breast cancer study [20], including gains in Chromosomes 1 and 8 and losses in Chromosome 4. As discussed in the Results, dependence on breast cancer molecular type is seen, but the sample size is to small to make the findings systematic.

Analysis of the OG/TSG ratio hypothesis ([43], Figure 2), which locates the high ratio values in increased copy number genomic regions, and the low ones in decreased copy number regions, confirms the hypothesis. The statistics are concurrent with the seminal paper by Watkins et al. [43] and a recent modeling study [13], and highly significant (see Results, Section 3.3). Our results indicate a significantly increased copy number in regions containing OGs and decreased in TSGs compared with normal ploidy of the sample. This outcome remain true if we assume as a reference the “normal” ploidy equal to 2, and also if we assume as a refrence the weighted average of all genomic regions in given sample). In addition, we found a significant difference in the distribution of copy number in regions encoding driver genes between primary tumor and lymph node samples, with the latter having generally higher copy number (on average and in regions containing driver genes).

We discussed at length in the Results section, how removal of spurious calls is changing the shape of the mutational spectra. To briefly summarize, the unfiltered spectra dominated by spurious FFPE-induced variant calls, are mostly right-skewed, which is only partly altered when the CCF spectra are computed (Figures 12 and 13). Filtering, particularly using DEEPOMICS software, brings out a finer structure of neutral and selective (“humplike”) components. They can be resolved in several specimens, with interesting conclusions drawn.

As noted before, according to the usual interpretation [44], for regularly diploid genomes, the centroid of the truncal hump should correspond to *f*_*h*_ = 0.5 *× ρ* frequency, where *ρ* is sample purity. This law is satisfied approximately in some case, while it is violated in other, as it can be concluded from examples in Section 3.4.2 (Figure 15). There exist at least three alternative interpretation of the apparently misplaced truncal humps: (i) Most of tumor mutations occur at non-diploid background (ii) purity is much worse in some cases than estimated by ASCAT, and (iii) the humps in the spectra are not truncal at all, but mixtures of secondary sub-clones of cells, as proposed in many sources, including [11]. Based on the current data, it is difficult to resolve this problem.

However, it is possible to analyze the inferred neutral components of the spectra in several cases, as we did in Section 3.4.2 (Table 3). Based on a simple theory developed in Section 2.2.4, we estimated the exponent *α* of the power law components of the P1 and L1 specimens, to obtain values consistently varying around *α* = 1. This led us to estimates 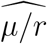 of the reduced mutation rates (mutation per division), which were consistently 10 to 100 times higher than in normal eukaryotic cells. Another interesting observation is generally high overlap of mutations in the primary tumors and lymph node metastases (Figure 14), which also applies to driver mutations that recur between the primary tumor and the lymph node (Fig. 8). This seems to support the hypothesis that metastasis and primary tumor are systematically exchanging cells, as opposed to metastases being seeded by a very limited subset of cells.

## 5 Conclusions

Summing up, despite the imperfections of the FFPE data, many important features of the molecular evolution of tumor DNA can be recovered from routine clinical samples. A positive association was found between the frequency of oncogenes relative to tumor suppressor genes and DNA copy number. In addition, interesting results concerning clonal structure, early tumor expansion, and interdependence of the primary tumor and lymph node metastases have been obtained.

## List of abbreviations

ABC-SMC-DRF: Approximate Bayesian Computation sequential Monte Carlo with random forests
BRCA: breast cancer
CCF: cancer cell fraction
CNV: copy number variation
EMT: epithelial-mesenchymal transition
ER: estrogene rceptor
FF: fresh frozen
FFPE: formalin-fixed, paraffin-embedded human epidermal growth factor receptor 2
HER2-: human epidermal growth factor receptor 2 negative breast cancer
HER2+: human epidermal growth factor receptor 2 positive breast cancer (HER2-enriched)
ISM: infinite-sites model
MRCA: most recent common ancestor
OG: oncogene
PR: progesterone receptor
SFS: site frequency spectrum
SNV: single-nucleotide variant
TNBC: triple-negative breast cancer
TSG: tumor suppressor gene
VAF: variant allele frequency
WGD: whole-genome duplication

## Declarations

### Ethics approval and consent to participate

The study received approval from the Ethical Committee at the Regional Medical Chamber in Krakow, Poland (decision of 4th December 2013). The permission does not receive a serial number, since it concerned de-identified retrospectively examined paraffin-embedded tumor specimens. Written informed consent for participation was not required for this study in accordance with the national legislation and the institutional requirements.

### Consent for publication

Not applicable.

### Availability of data and materials

Breast cancer sequencing data can be found at https://ega-archive.org/ with accession number: EGAD00001009081. Any queries should be directed to the corresponding author.

### Authors’ contributions

Conceptualization, M.K., R.J. and M.K.K.; methodology, M.K., R.J.; sample collection and preparation, A.A. and K.M.; software, M.K.K., P.K., K.N.D and R.J.; validation, M.K.K. and M.K.; formal analysis, M.K.; investigation, M.K.K., P.K., K.N.D. and M.K.; resources, M.K.K., R.J. and P.K.; data curation, M.K.K., R.J. and P.K.; writing—original draft preparation, M.K.K. and M.K.; writing—review and editing, M.K.K. and M.K.; visualization, M.K.K., P.K and K.N.D.; supervision, M.K.; project administration, M.K.; funding acquisition, M.K. All authors read and approved the final manuscript.

## Acknowledgments

We would like to thank Inyoung Kim from Theragen Bio for conducting the DEEPOMICS and DEEPOMICS FFPE Plus analyses using the on-premise version of both tools. The results shown here are in part based upon data generated by the TCGA Research Network: https://www.cancer.gov/about-nci/organization/ccg/research/structural-genomics/tcga. Computations were carried out using the infrastructure of the Ziemowit computer cluster (www.ziemowit.hpc.polsl.pl) in the Laboratory of Bioinformatics and Computational Biology, The Biotechnology, Bioengineering and Bioinformatics Centre Silesian BIO-FARMA, created in the POIG.02.01.00-00-166/08 and expanded in the POIG.02.03.01-00-040/13 projects.

## A Appendix: Data acquisition and cleaning

### A.1 Additional files

Supplementary Table 1: Sequencing statistics for all BRCA samples (file: Supplementary Table 1.xlsx; 18 KB)

### A.2 DNA sequencing quality control

Shown below are the exon coverage statistics for all samples in the cohort (Figure S1). The sequencing statistics are presented in Supplementary Table 1. They list the read statistics for all samples, subdivided into primary tumor (P1), cancerous lymph node (L1) and benign lymph node (C), used as control for the detection of somatic mutations.

The read statistics for our BRCA group are shown in Figure S2. The graph showing sequencing depth (coverage) is included as Figure S3.

**Figure S1:**
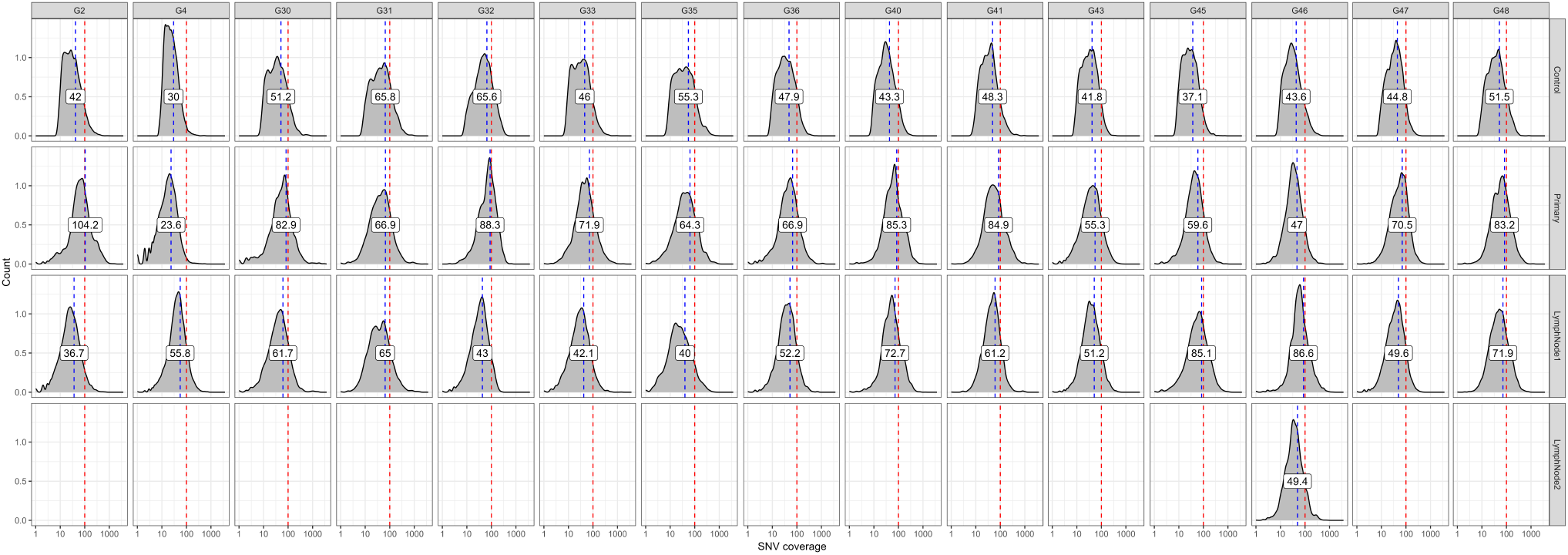
Exon coverage characteristics for all BRCA samples.

**Figure S2:**
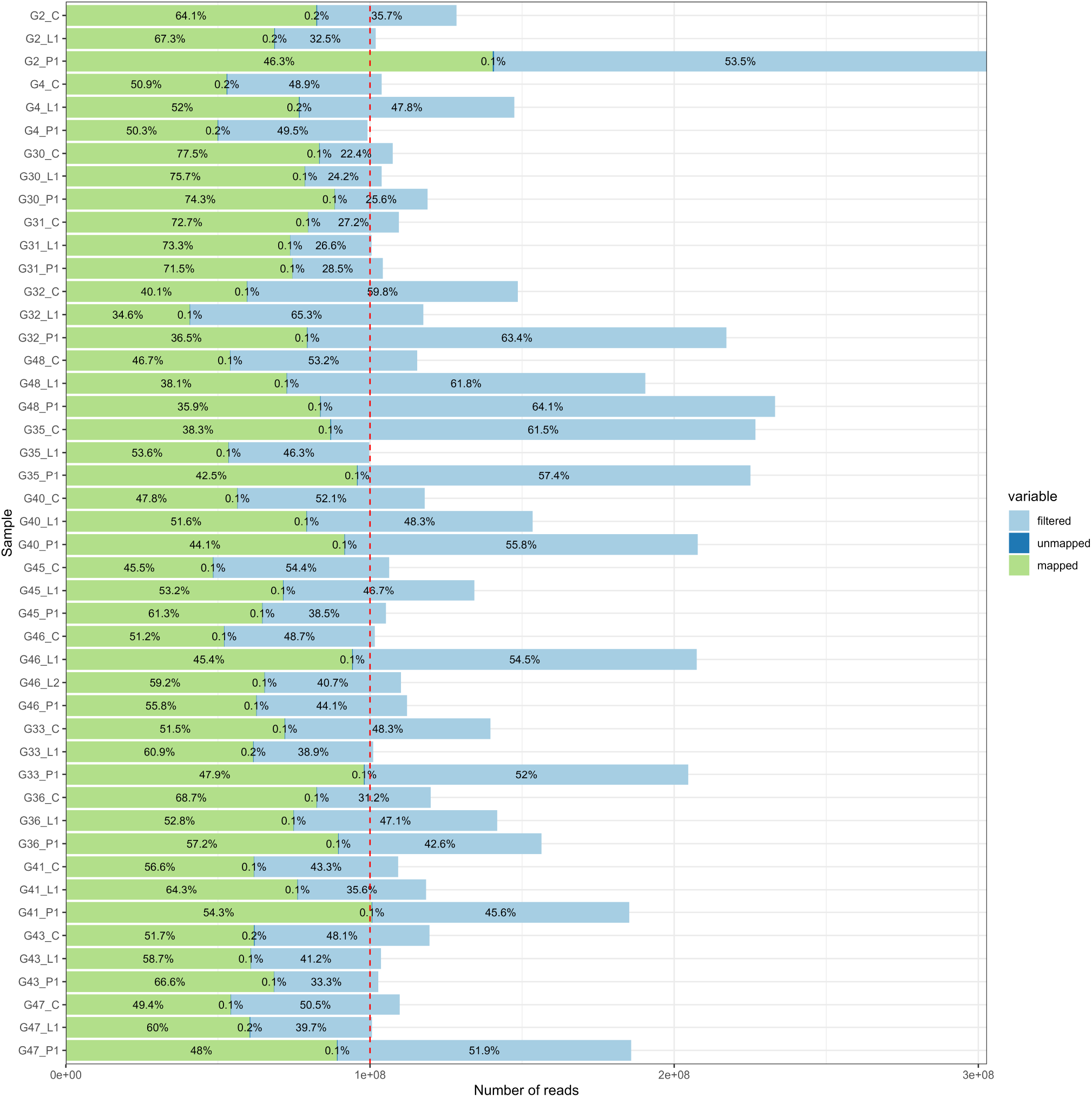
Read statistics for all BRCA samples.

**Figure S3:**
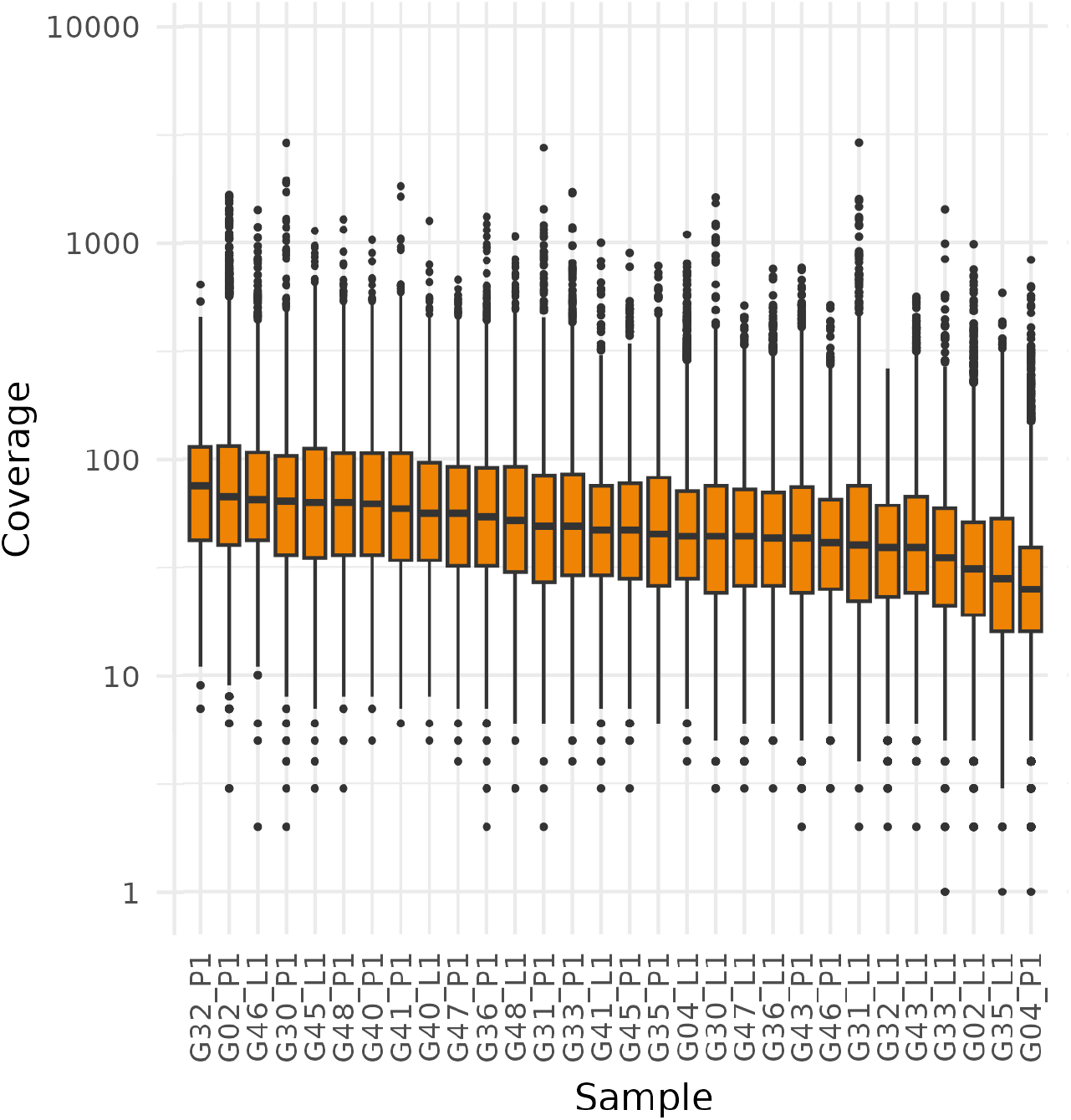
Sequencing coverage of SNVs and Indels across samples in BRCA cohort.

Figure S4 shows the count of variants retained in each sample after filtration with each tool and, additionally, the count of variants in the case when FPPE score threshold was changed to 0.25 in the result of DEEPOMICS. The numbers were highest in case of SOBDetector, significantly exceeding these observed after filtration with other tools and also the average number of variants observed in TCGA samples. In turn, the counts of variants obtained with other tools were too low for some of the analyses we conducted, and therefore we decided to alter the FFPE score threshold in DEEPOMICS to enrich the outcome with variants with lower allele frequency, but retaining sufficient efficacy of filtration (see Figure 5).

**Figure S4:**
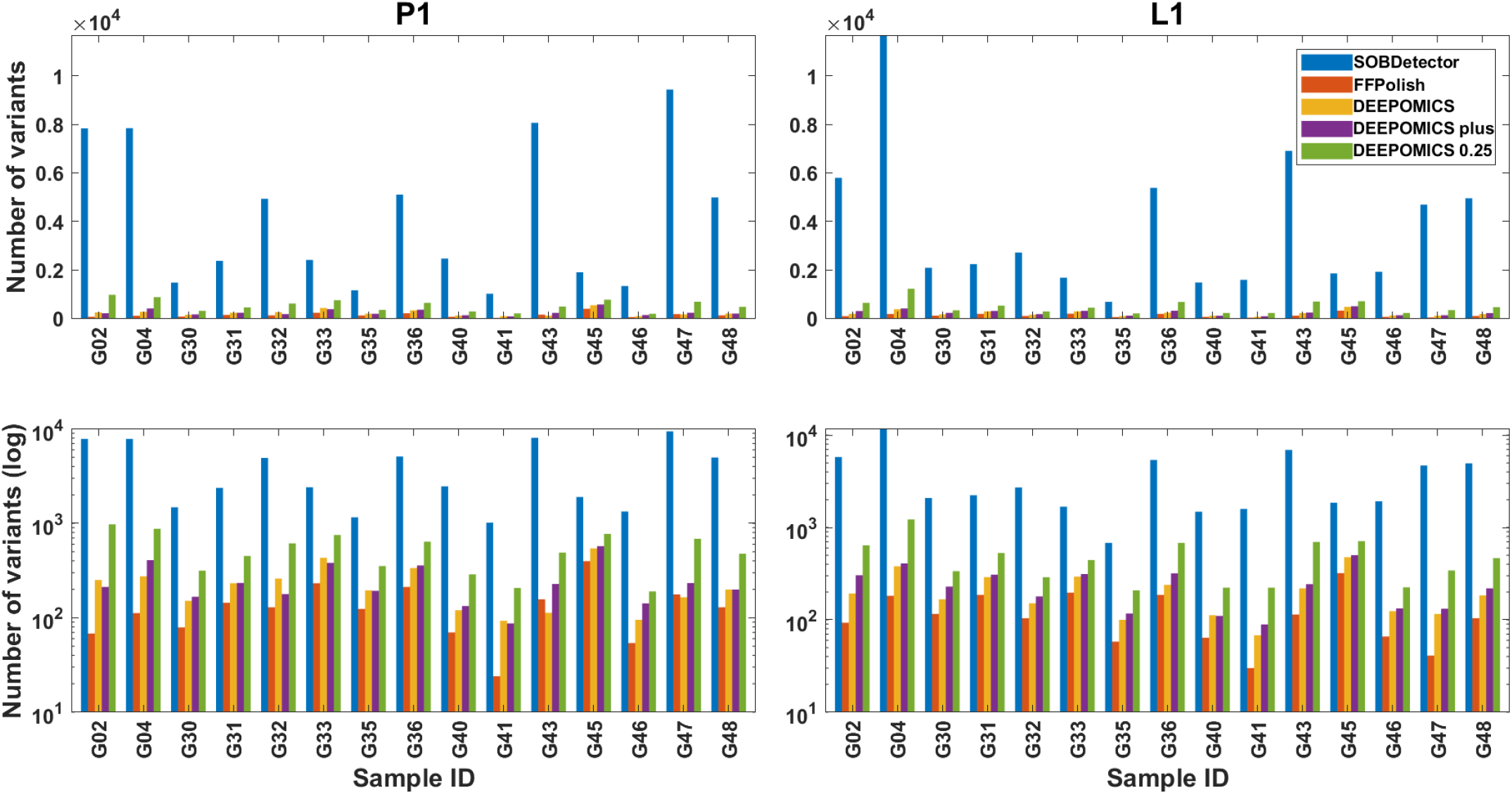
Count of single nucleotide variants retained in primary tumor (P1, left column) and lymph node metastasis (L1, right column) samples by each of the tools and number of variants for DEEPOMICS with adjusted FFPE score threshold (DEEPOMICS 0.25). Both sample groups are shown in linear (top row) and log (bottom row) scale.

## B Appendix: Additional details of genomic analysis

### B.1 Driver ploidy

Figure S5 shows the result of analysis of copy number change in genomic regions encoding driver mutations in all samples before (top panel) and after filtration with DEEPOMICS with adjusted FFPE score threshold (bottom panel). The background dashes (horizontal for P1 and vertical for L1) represent relative copy number in region with respect to assumed normal ploidy of sample equal to weighted average of all genomic regions in this sample. The plot area of both figures is divided into three major groups: tumor suppressor genes (TSG), oncogenes (OG) and a class of genes (OG/TSG) acting in different ways depending on specific context, such as e.g. a BRCA subtype or a specific mutation site.

For such estimated baseline ploidy, which in most cases is higher than 2, the proportion of copy number gains (marked with red color in the graph) is lower than for reference ploidy equal to 2. Instead, we observe the decrease of copy number in regions containing tumor suppressor genes. The region containing TP53 gene (here classified as a mixed-role gene) is characterized by lower than baseline copy number and this effect is shared by most of the samples which have an additional mutational hit in this gene.

**Figure S5:**
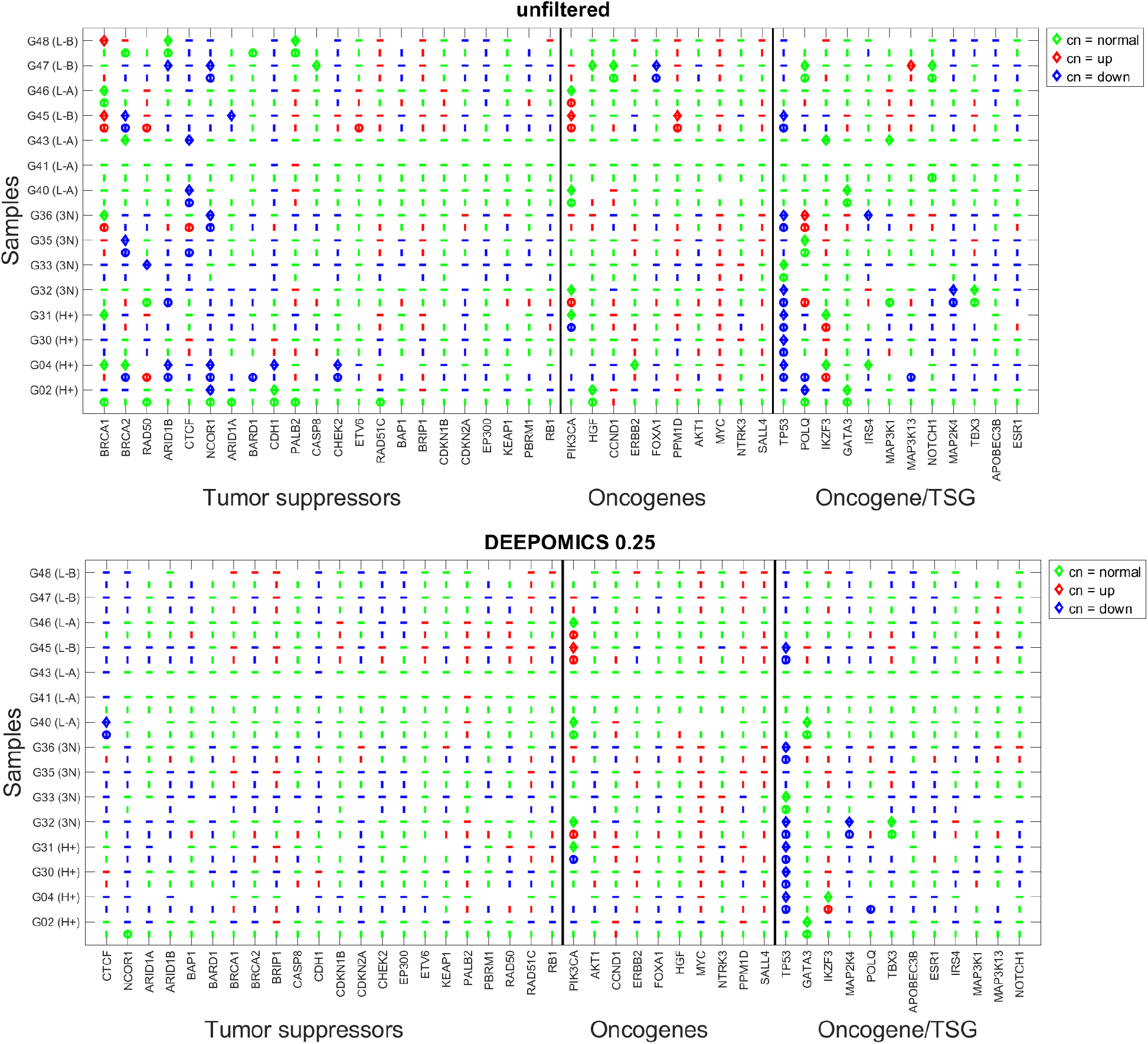
Ploidy in genomic regions encoding driver mutations in all P1 and L1 samples before (top panel) and after filtration with DEEPOMICS with adjusted FFPE score threshold (bottom panel). Dashes (horizontal for P1 and vertical for L1) represent relative ploidy in region with respect to assumed normal ploidy of the sample equal to weighted average of all genomic regions in this sample. The change in ploidy is encoded with color: green - normal ploidy, red - increased ploidy, blue - decreased ploidy. Additionally mutations in driver genes are marked by diamond (P1) and circle (L1). Horizontal axis of the graph is sorted by driver gene function and divided into 3 major groups: TSG’s, OG’s and OG/TSG’s. In each of the major groups driver genes are sorted in descending number of mutational hits.

**Figure S6:**
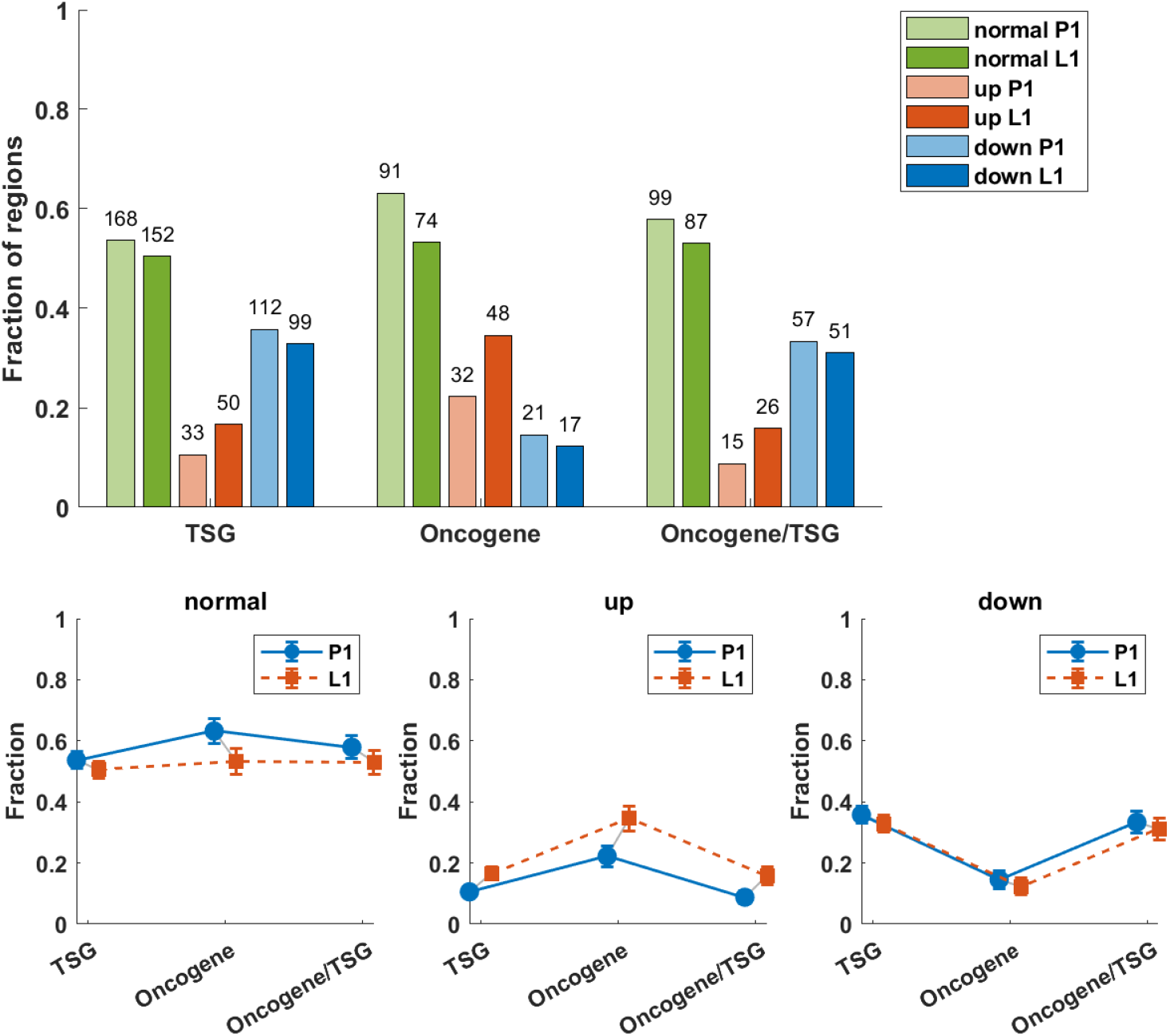
Statistics of copy number status in regions encoding driver mutations across all P1 and L1 samples, assuming normal ploidy of the sample equal to weighted average of all genomic regions in this sample. **Top panel:** Average copy number in regions encoding three major groups of driver genes: TSG, OG, and OG/TSG. **Middle panel:** Fractions of regions with copy number increased (red), decreased (blue) or not changed (green) with respect to assumed normal ploidy equal to 2. The height of the bar depicts proportional share in given major group, while the numbers above are the counts of regions of given type. **Bottom panel:** Fractions of regions encoding driver mutations in P1 and L1 with copy number not changed (normal), increased (up) and decreased (down) in the three major groups of driver genes. Distribution of TSG and OG in P1, comparing copy number down vs. normal vs. up (counts: down — TSG 112, OG 21; normal — TSG 168, OG 91; up — TSG 33, OG 32): *χ*^2^ = 26.27, *p <* 0.001. In L1 (counts: down — TSG 99, OG 17; normal — TSG 152, OG 74; up — TSG 50, OG 48): *χ*^2^ = 29.25, *p <* 0.001. Both results are significant at *p <* 0.05 — TSGs are enriched with copy number loss and OGs are enriched among amplifications. Total counts of regions with copy number down/normal/up (TSG, OG, OG/TSG combined) across P1 and L1 (counts: down — P1 190, L1 167; normal — P1 358, L1 313; up — P1 80, L1 124): *χ*^2^ = 13.53, *p <* 0.001. The result is significant at *p <* 0.05 — P1 shows relatively more copy number losses while L1 shows relatively more amplifications.

We checked how many BRCA-driver containing genomic regions fall in the category of increased, decreased and normal (unchanged) copy number. Results are subdivided into three groups: TSG, OG and OG/TSG. In all three groups fraction of regions with unchanged copy number is higher than in the remaining categories.

The fraction of regions with increased and decreased copy number is similar between TSG and OG/TSG groups while in the OG group, the highest increase in copy number is observed. In addition, the fraction of genomic regions with decreased copy number is the smallest.

This trend, which is most clearly shown by Figure 11 remains present also in the case of estimated baseline ploidy of the samples. All 3 *×* 2 comparisons were tested using two-tailed *χ*^2^ test for contingency tables of gene class (TSG vs OG) or sample (P1 vs L1) versus outcome (down vs normal vs up). Counts are the numbers of genomic regions.

### B.2 Site frequency spectra

In this section, we first present aggregates of all samples. In the aggregate spectra as depicted in Figure S7, shown are also the positions of BRCA drivers in the frequency spectrum. The counts of driver genes at particular VAF frequencies are marked with red and shown in the logarithmic scale.

To take into account purity and ploidy which vary across samples we calculated Cancer Cell Fraction (see Methods, Section 2.2.2) and presented it in the same form as the aggregated VAF-based site frequency spectrum (Figure S8). As in the previous case, the bars at frequencies where driver mutations were found are marked with red.

In both cases, in unfiltered and SOBDetector processed datasets driver mutations are present in almost all lowest frequency-bars (above the 0.05 threshold for VAF imposed on mutations in regions containing driver genes), while in a growing and mutationg tumor they are expected to be shared by a larger proportion of variant calls. For the FFPolish- and DEEPOMICS-filtered data most drivers fall into frequency thresholds from 0.2 to 0.5, which after correction for purity and ploidy and calculation of CCF spectrum shifts them towards frequency close to 1.0. Please see the cluster of drivers around value 1 in panels depicting CCF spectra under FFPolish and DEEPOMICS filtrations. In additon, see Equ. (7), which explains the relationship between the scales of the CCF-vs. VAF-based spectra.

**Figure S7:**
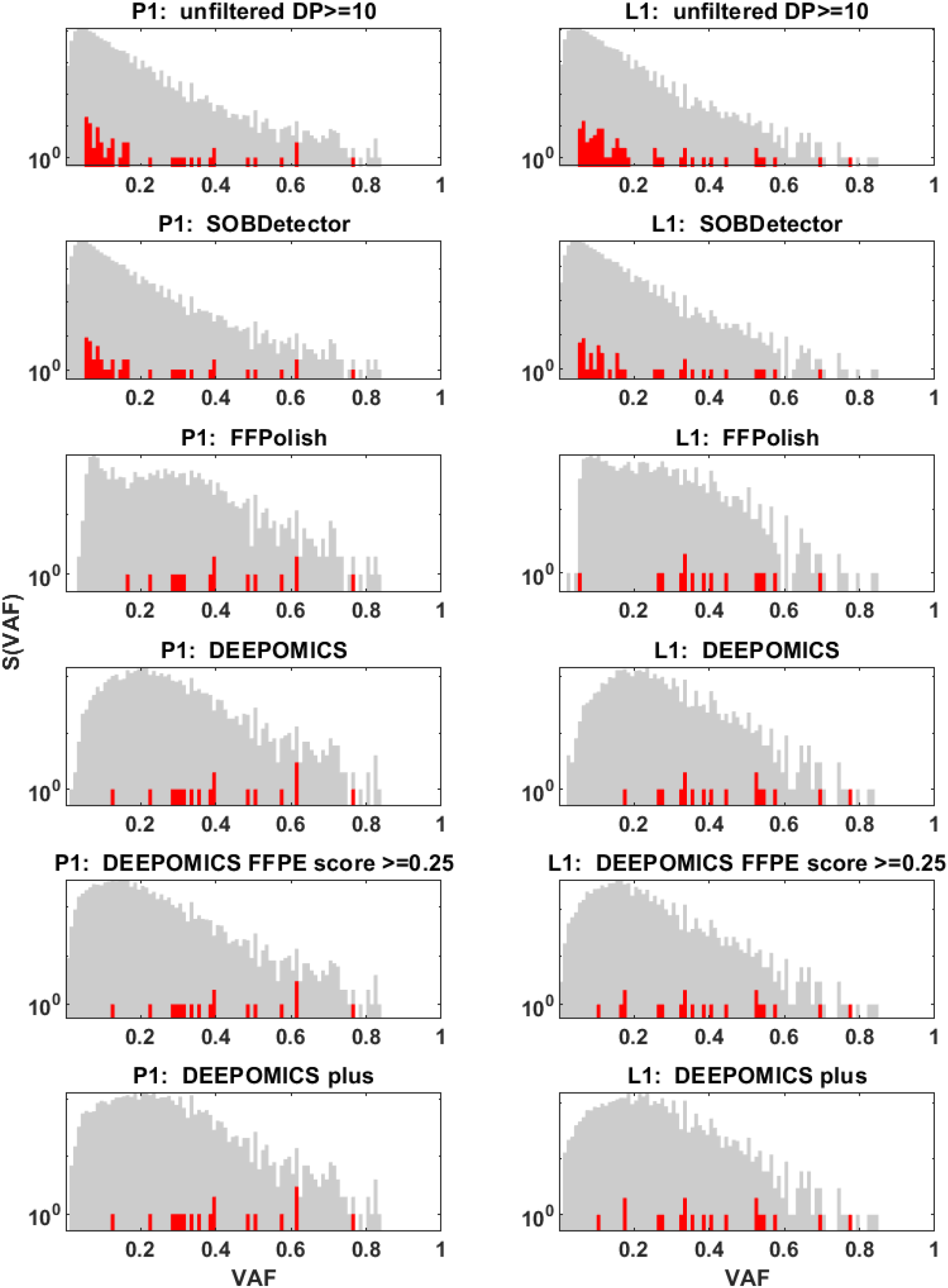
Site frequency spectrum (SFS) based on aggregated variant allele frequency (VAF) from patients samples from primary tumor (P1, left column) and lymph node metastasis (L1, right column) in the log scale. With red marked are bars for VAF thresholds in which mutation variants in driver genes were observed with actual number of drivers belonging to this threshold. Each panel from top to bottom was generated using different filtration method. Notice the differences on Y axis between methods.

**Figure S8:**
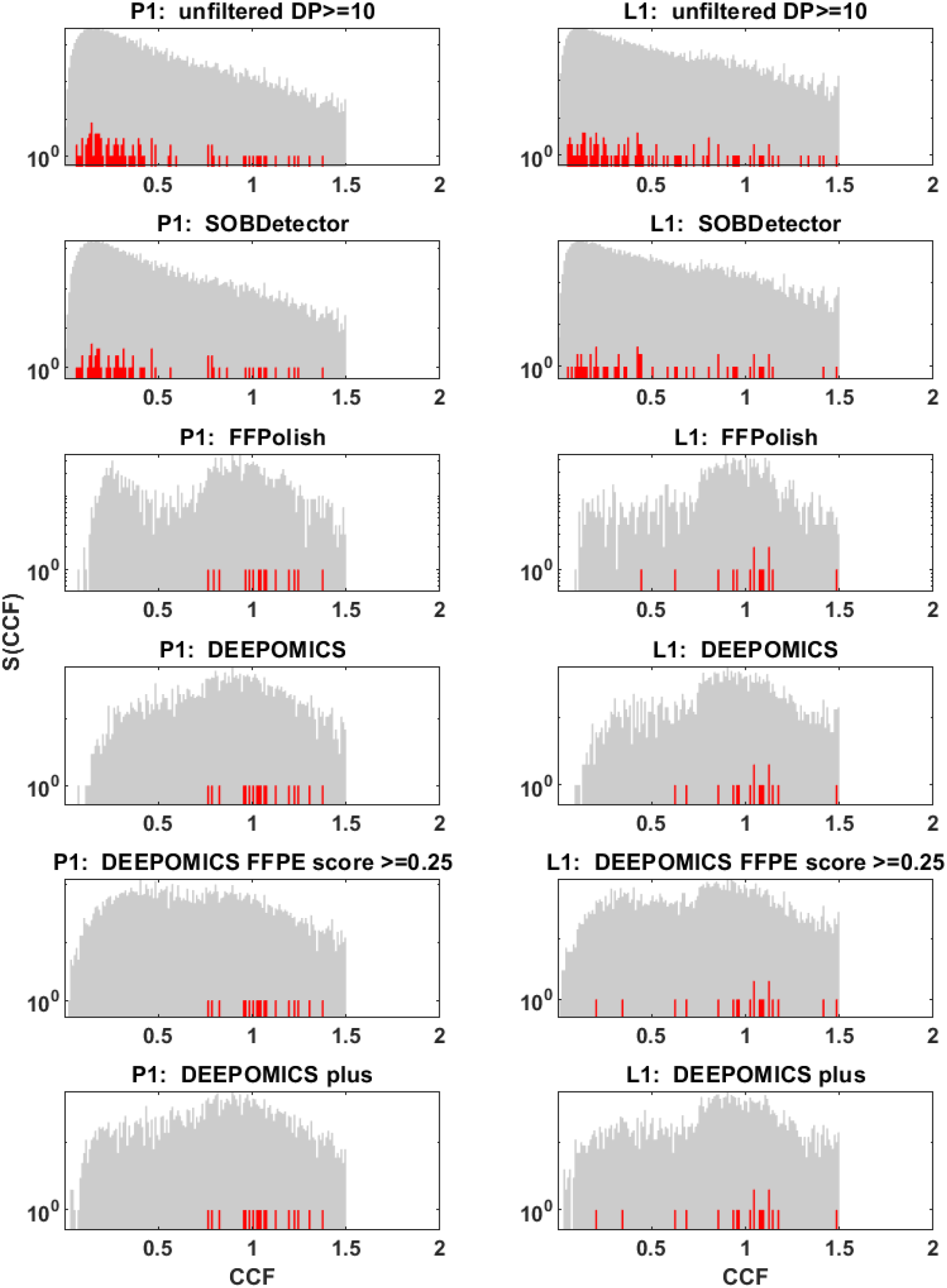
Site frequency spectrum (SFS) based on aggregated cancer cell fraction (CCF) from patients samples from primary tumor (P1, left column) and lymph node metastasis (L1, right column) in the log scale. With red marked are bars for VAF thresholds in which mutation variants in driver genes were observed with actual number of drivers belonging to this threshold. Each panel from top to bottom was generated using different filtration method. Notice the differences on Y axis between methods and the range of X scale (CCF not rescaled).

#### B.2.1 Listing of SFS and CCF spectra of all specimens

Figures S9-S12 represent site frequency spectra for individual samples, based on VAF (Figs S9 and S10) and CCF (Figs S11 and S12). Shown are results before (Figs S9 and S11) and after filtration (Figs S10 and S12).

**Figure S9:**
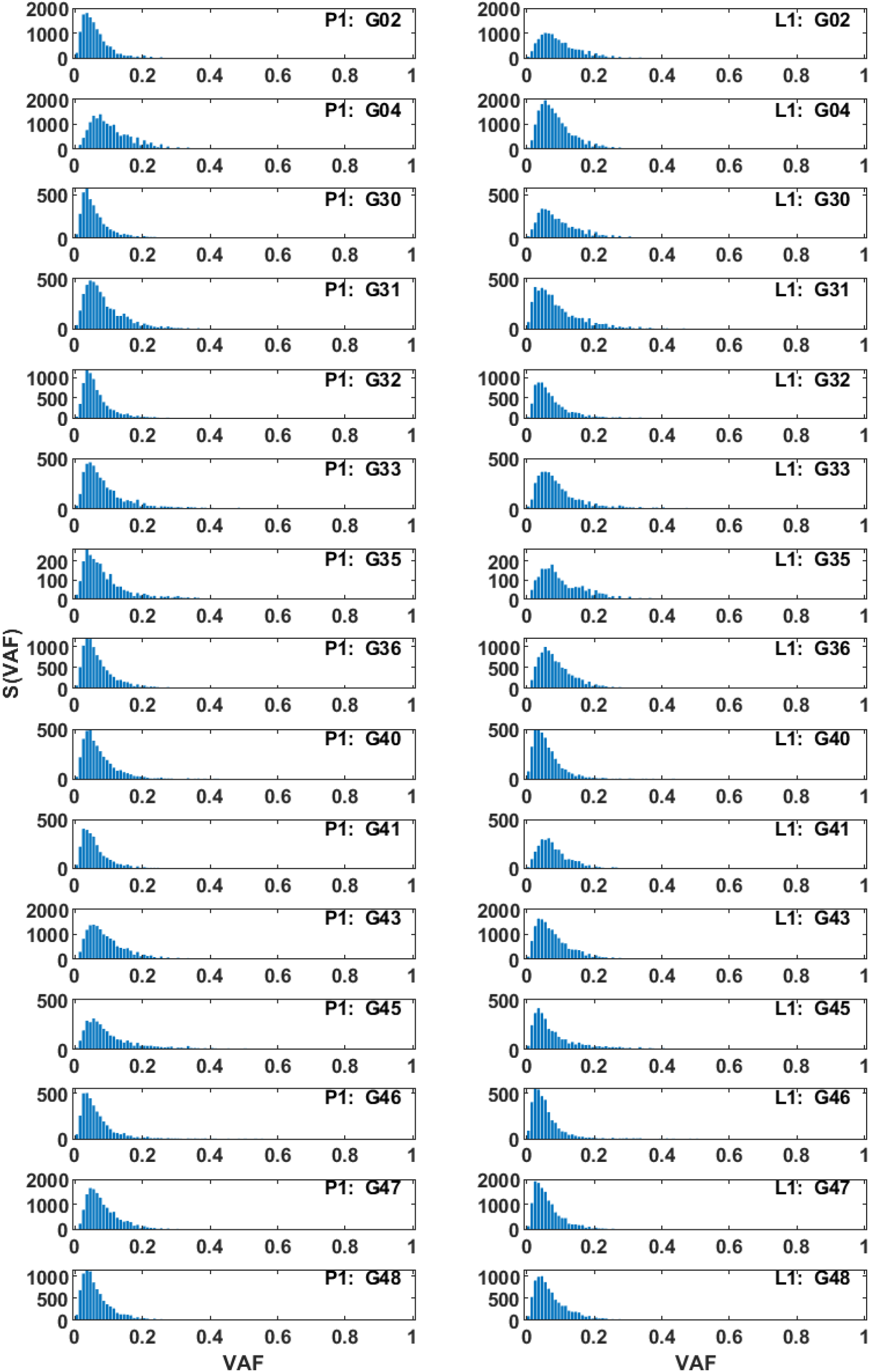
SFS based on VAF for all samples (unfiltered) for primary tumor (P1, left column) and lymph node metastasis (L1, right column).

**Figure S10:**
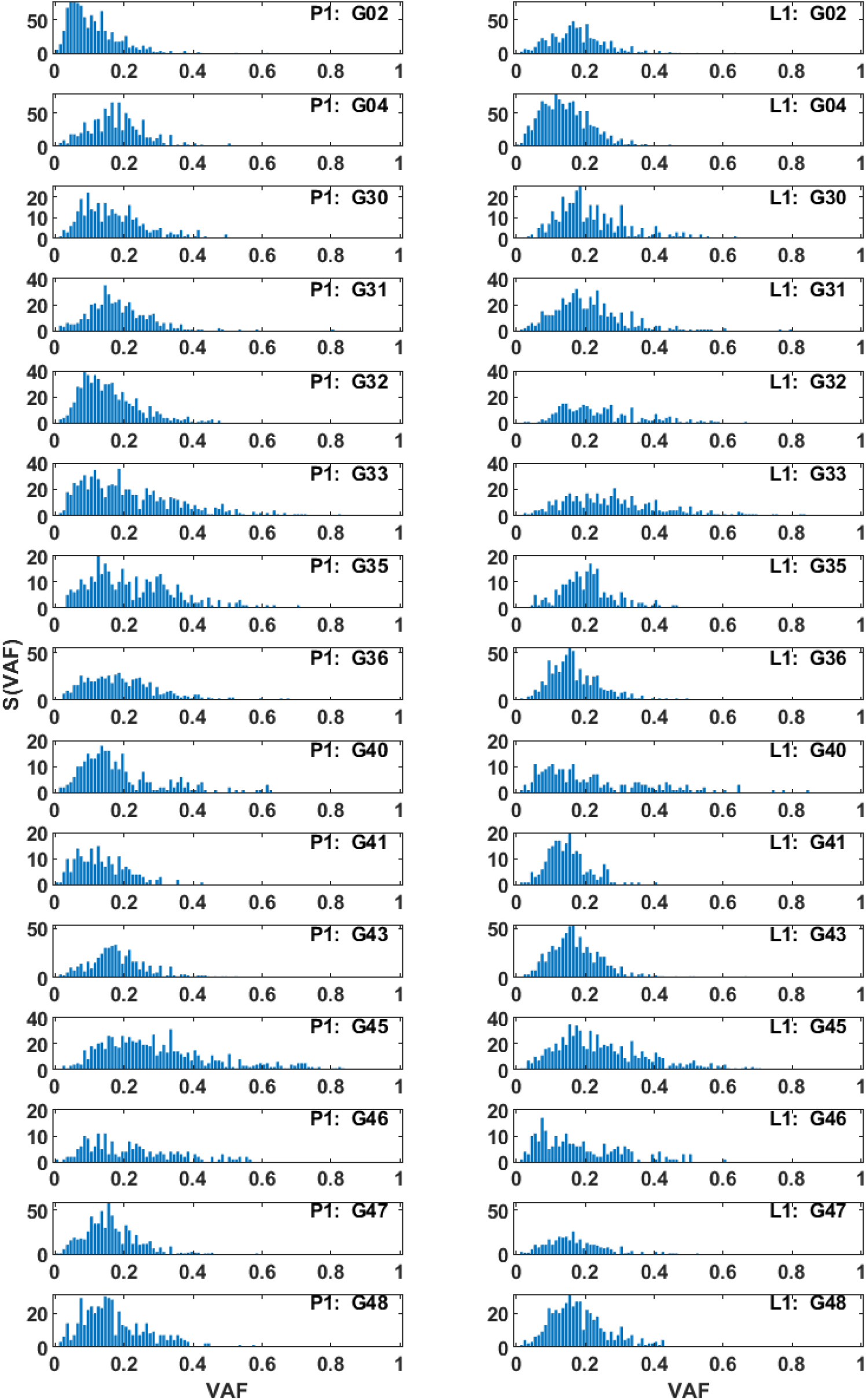
SFS based on VAF for all samples (filtered with DEEPOMICS with FFPE score threshold= 0.25) for primary tumor (P1, left column) and lymph node metastasis (L1, right column).

**Figure S11:**
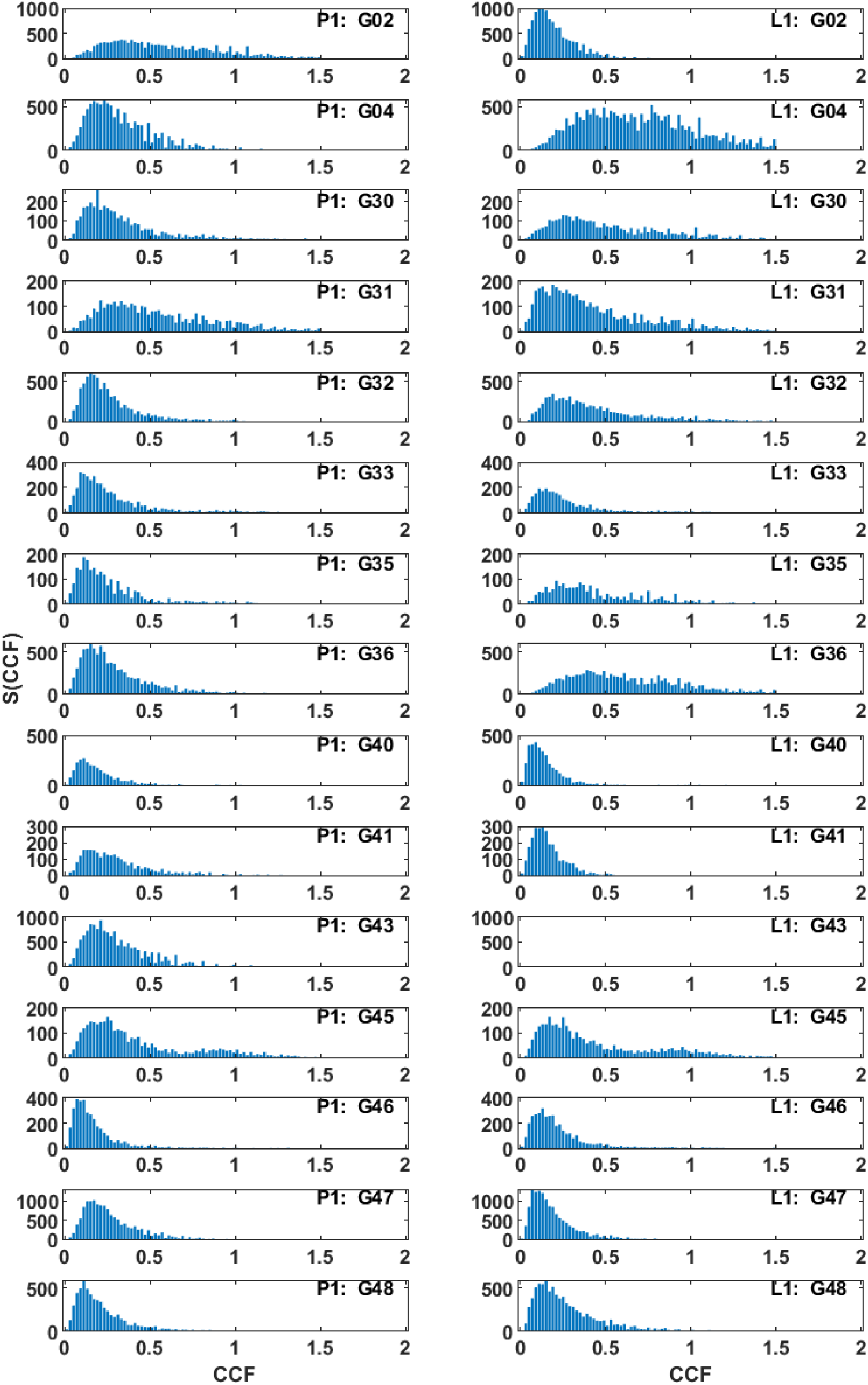
SFS based on CCF for all samples (unfiltered) for primary tumor (P1, left column) and lymph node metastasis (L1, right column).

**Figure S12:**
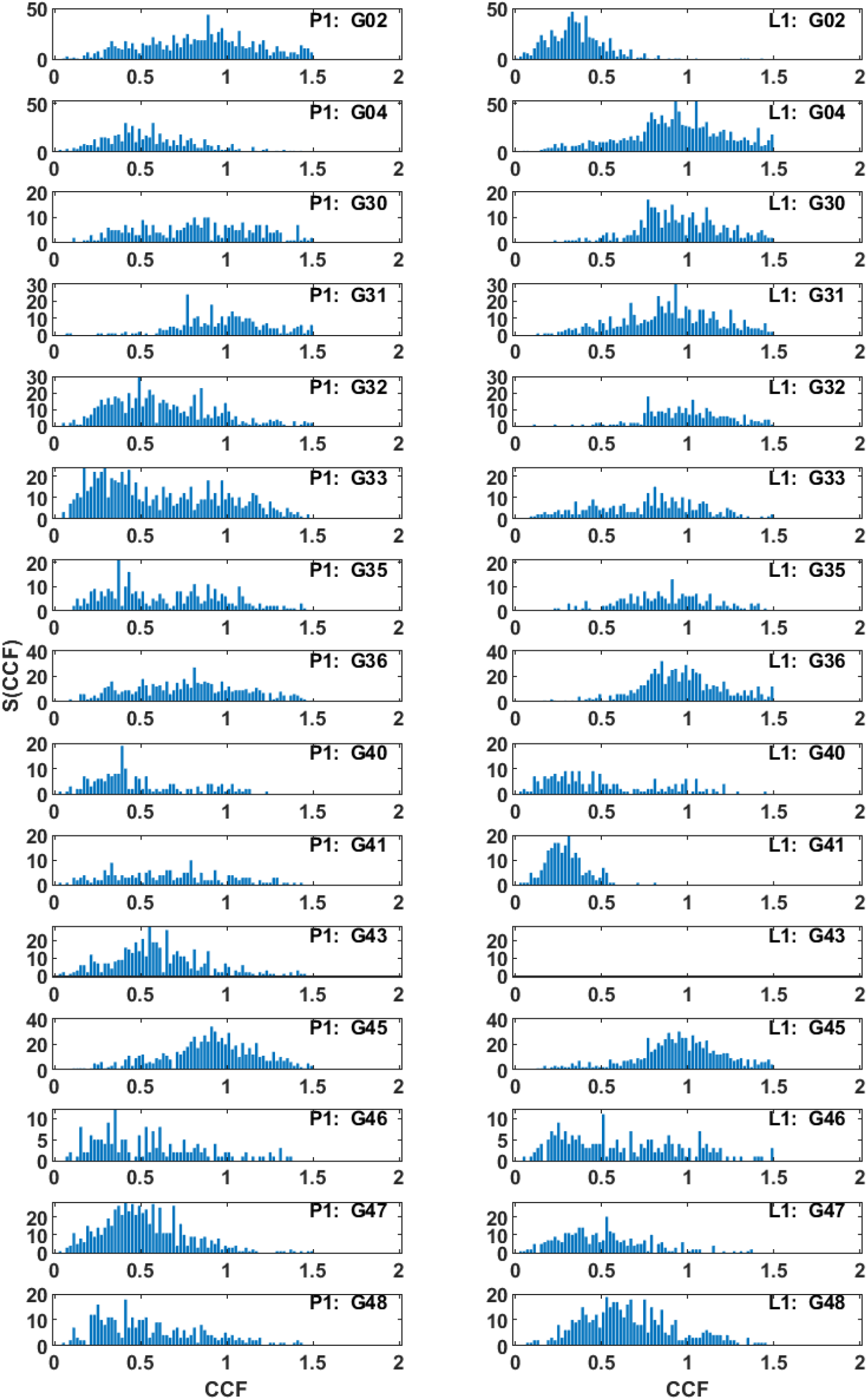
SFS based on CCF for all samples (filtered with DEEPOMICS with FFPE score equal to 0.25) for primary tumor (P1, left column) and lymph node metastasis (L1, right column).

### B.3 Trade-off between purity and ploidy estimates

For each purity-ploidy pair ASCAT [42] rounds the inferred tumor copy numbers to integers and computes how well these integers determine the observed LogR and BAF [39]. Different integer assignments yield locally good fits, producing disjoint curved bands (valleys) of low error in which the segments can be explained by the same integer copy number combination (Fig. S13). In some cases, the result obtained might be ambiguous due to trade-off between purity and ploidy estimates, which results in multiple regions with low error.

Copy number plots (Fig. S14) show pointwise SNP data across the genome with two rows: the per-locus LogR (log_2_ total intensity ratio) and the BAF (B-allele fraction). LogR reports changes in total DNA amount while BAF reports allelic imbalance — ASCAT fits both jointly (under a tumor+normal mixture and integer tumor copy numbers) to infer purity, ploidy and allele-specific copy number.

**Figure S13:**
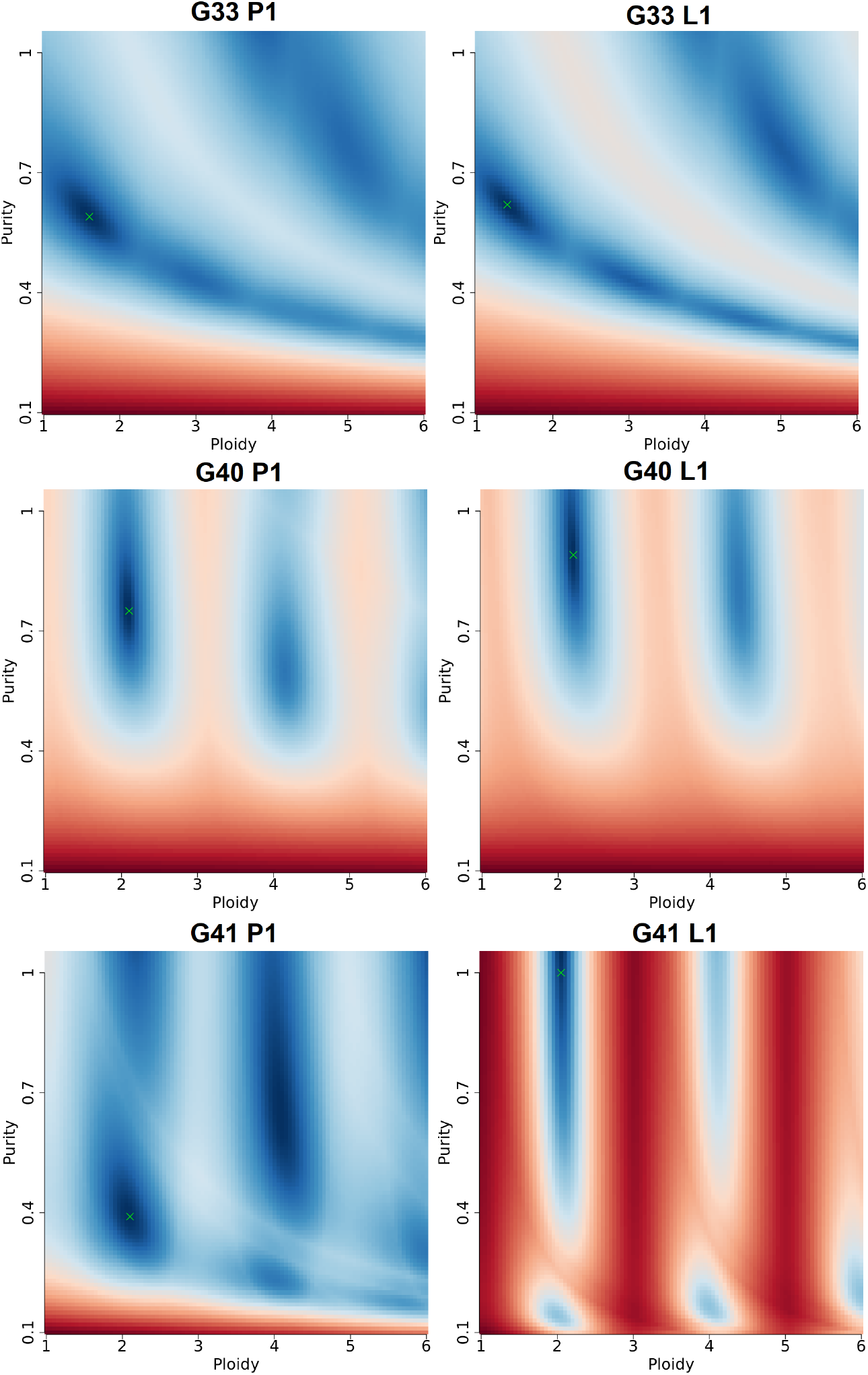
Ploidy-purity plots for patients G33, G40 and G41 for primary tumor (P1, left column) and lymph node metastasis (L1, right column). Sunrise plots represent a fit-score surface over a grid of candidate purity (y-axis) and ploidy (x-axis) values. Color encodes goodness of fit: dark blue regions represent good fit (low error), red regions correspond to worse fit. The green crosses mark the ASCAT-selected solutions (the ploidy-purity pairs chosen for downstream integer copy-number calling).

**Figure S14:**
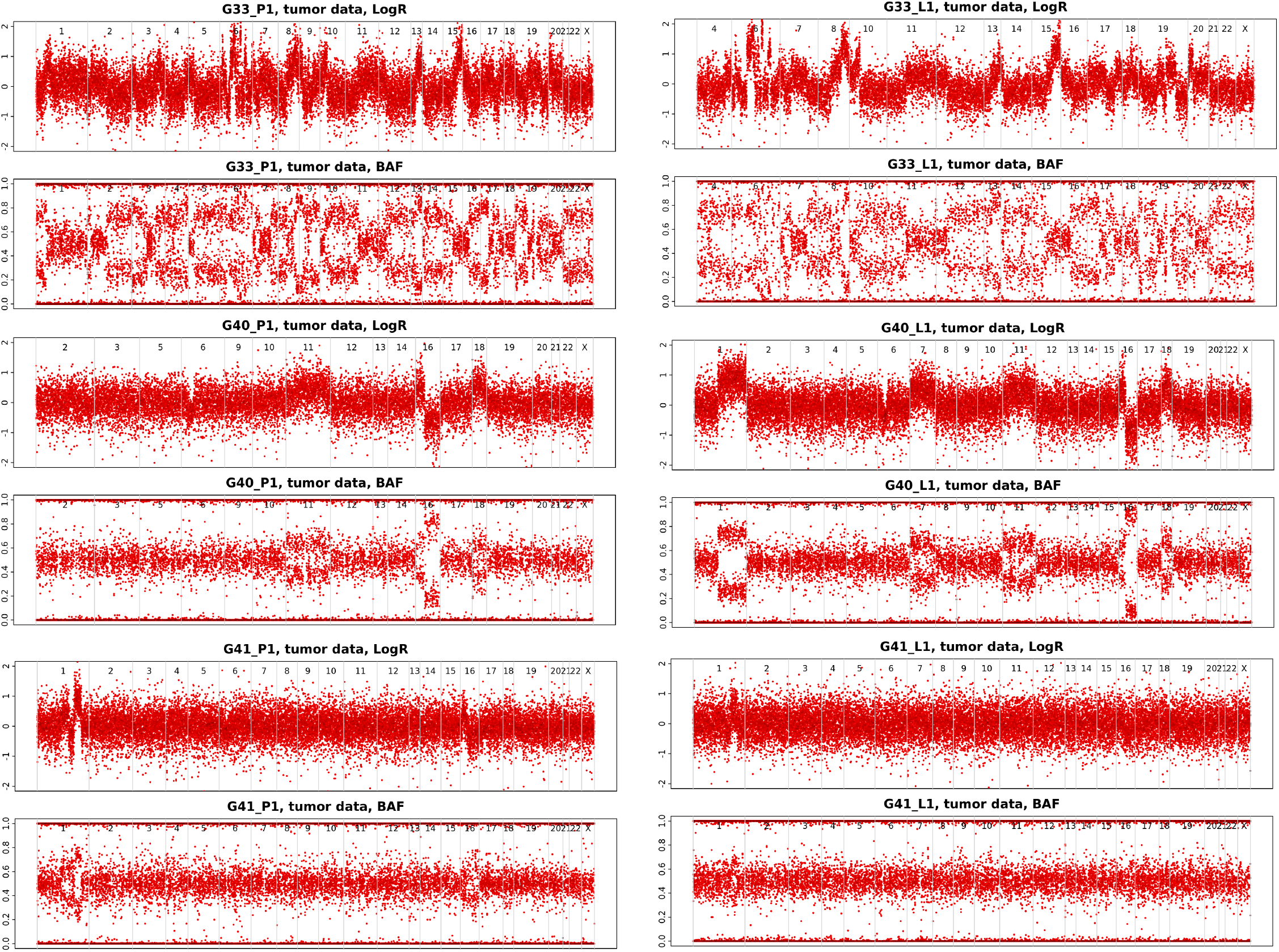
Copy number plots after GC and replication-timing (RT) correction for patients G33, G40 and G41 for primary tumor (P1, left column) and lymph node metastasis (L1, right column) show pointwise SNP data across the genome in the form of the per-locus LogR (top panel) and the BAF (bottom panel). Vertical separators mark chromosomes.

